# Brown adipocytes local response to thyroid hormone is required for adaptive thermogenesis in adult male mice

**DOI:** 10.1101/2022.08.03.502639

**Authors:** Yanis Zekri, Romain Guyot, Inés Garteizgogeascoa Suñer, Laurence Canaple, Amandine Gautier-Stein, Justine Vily-Petit, Denise Aubert, Sabine Richard, Frédéric Flamant, Karine Gauthier

## Abstract

Thyroid hormone (T3) and its nuclear receptors (TR) are important regulators of energy expenditure and adaptive thermogenesis, notably through their action in the brown adipose tissue (BAT). However, T3 acts in many other peripheral and central tissues which are also involved in energy expenditure. The general picture of how T3 regulates BAT thermogenesis is currently not fully established, notably due to the absence of extensive omics analyses and the lack of specific mice model. Here, we first used transcriptome and cistrome analyses to establish the list of T3/TR direct target genes in brown adipocytes. We then developed a novel model of transgenic mice, in which T3-signaling is specifically suppressed in brown adipocytes at adult stage. We addressed the capacity of these mice to mount a thermogenic response when challenged by either a cold exposure or a high-fat diet, and analyzed the associated changes in BAT transcriptome. We conclude that T3 plays a crucial role in the thermogenic response of the BAT, controlling the expression of genes involved in lipid and glucose metabolism and regulating BAT proliferation. The resulting picture provides an unprecedented view on the pathways by which T3 activates energy expenditure through an efficient adaptive thermogenesis in the BAT.

**Significance Statement:** Thyroid hormones (TH) increase energy expenditure by regulating the expression of target genes in many metabolic tissues. Among them, brown adipose tissue (BAT) dissipates biochemical energy into heat production to notably prevent hypothermia during cold exposure. Hypothyroid mice display inefficient BAT thermogenesis suggesting that TH are crucial for this process. Here, we eliminated TH signaling specifically in brown adipocytes and expose the mice to different physiological stressors. We showed that TH signaling is crucial for BAT thermogenesis as it controls the expression of genes involved in proliferation and in the metabolism of lipids and glucose, the main energy resources for BAT thermogenesis. This study provides an unprecedented view on the pathways by which T3 activates energy expenditure the BAT.

## Introduction

Brown adipose tissue (BAT) has the ability to dissipate energy through thermogenesis in response to cold and high-fat diet, to prevent hypothermia and limit body weight gain, respectively. In response to these stressors, sympathetic nerves stimulate brown adipocyte’s adrenergic receptors to trigger the cAMP-protein kinase A (PKA) ^1^, favoring local lipolysis and glycolysis to fuel an increase in energy metabolism ^2^. This is used by the BAT-specific protein UCP1 to favor thermogenesis at the expense of ATP production ^3^. In the long run, this causes an increase in lipogenesis to refill local lipid stocks, mitochondrial biogenesis and a proliferation of brown adipocytes ^4–6^. These actions are coordinated by transcription factors, notably involving the coactivator PGC1α ^7^. Importantly, cold exposure induces an indispensable increase expression and activity of the type 2 deiodinase activity (DIO2) resulting in an intracellular conversion of the inactive form of thyroid hormone, thyroxine or T4, into its active form, 3,3’,5-triiodo-L-thyronine or T3 ^4,8,9^.

T3 exerts a large influence on energy expenditure, as hypothyroidism is associated with cold sensitivity and weight gain, while hyperthyroid patients display the opposite phenotype (9,10,11). T3 binds to the broadly-expressed thyroid hormone nuclear receptors, TRα1 and TRβ1/2 (collectively called TR) encoded by the *Thra* and *Thrb* genes ^13^. Majority of TR bind to specific response elements resembling the archetypical DR4 consensus sequence (5’AGGTCAnnnnRGGnCA3’) to repress expression of neighboring genes. Upon T3 binding, TR results in a rapid activation of transcription ^14^.

T3 concentration drastical increase in BAT during cold exposure ^15^ is crucial, as *DIO2KO* mice have impaired BAT thermogenesis ^16^, notably due to altered fatty acids oxidation ^4^. However, the involvement of T3 in energy metabolism is not restricted to brown adipocytes, as it directly increases hepatic lipid and glucose metabolism ^17^, and favors exercise-associated thermogenesis in the muscle ^18^. It also triggers white adipose tissue (WAT) browning, a process in which heat-producing “beige” adipocytes expressing UCP1 emerge in the WAT ^19,20^. Finally, T3 stimulates indirectly BAT thermogenesis by acting in the hypothalamic ventro-medial nuclei where it triggers the sympathetic outflow to BAT ^21^.

Thus, T3 exerts a broad influence on different tissues to regulate energy expenditure, making the relative importance of BAT thermogenesis in T3-dependent energy expenditure unclear. Moreover, T3 action directly in the BAT has not been exhaustively described as, 1) no transcriptomic and cistromic analyses have been led to obtain T3 target genes in this tissue, 2) T3 also indirectly controls the BAT through the brain, making the role of local T3 difficult to pinpoint. Here, we aimed at distinguishing these two levels and focus on the T3 cell-autonomous (i.e., local) function in brown adipocytes, and address its importance in adaptive thermogenesis. To address this aim, we combined genomic analysis and the phenotyping of mice with selective blockade of T3-signaling in brown adipocytes. The resulting picture provides an unprecedented view on the pathways by which T3 activates energy expenditure through an efficient adaptive thermogenesis in the BAT.

## Results

### BATKO mice present a BAT-specific deletion of T3 signaling

The blockade of T3 signaling specifically in brown adipocytes has been achieved using the *Ucp1-CreER^T2^* (Suppl. figure 1). It was combined with two available floxed alleles to ascertain a full blockade of the T3 response of brown adipocytes. In the absence of tools to eliminate *Thra*, we used *Thra^AMI^*, a recombinant “floxed” allele of the *Thra* gene, in which Cre-mediated recombination allows the expression of TRα1^L400R^, a dominant negative version of the TRα1 receptor ^22^. TRα1^L400R^ prevents the recruitment of coactivators and is thus condemned to constitutively repress T3 target genes expression. Thus, we expect the effects of this knock-in to be stronger than a knock-out. Then, we used *Thrb^lox^*, a recombinant allele of *Thrb* gene, in which two loxP sequences flank the exon encoding the DNA binding domain of TRβ1/TRβ2 ^23^. This approach eliminates T3 responsiveness. *Ucp1-CreER^T2^xThra^AMI/+^Thrb^lox/lox^* mice (Figure 1A) are called BATKO mice and compared with *Thra^AMI/+^Thrb^lox/lox^* CTRL littermates.

**Figure 1:**
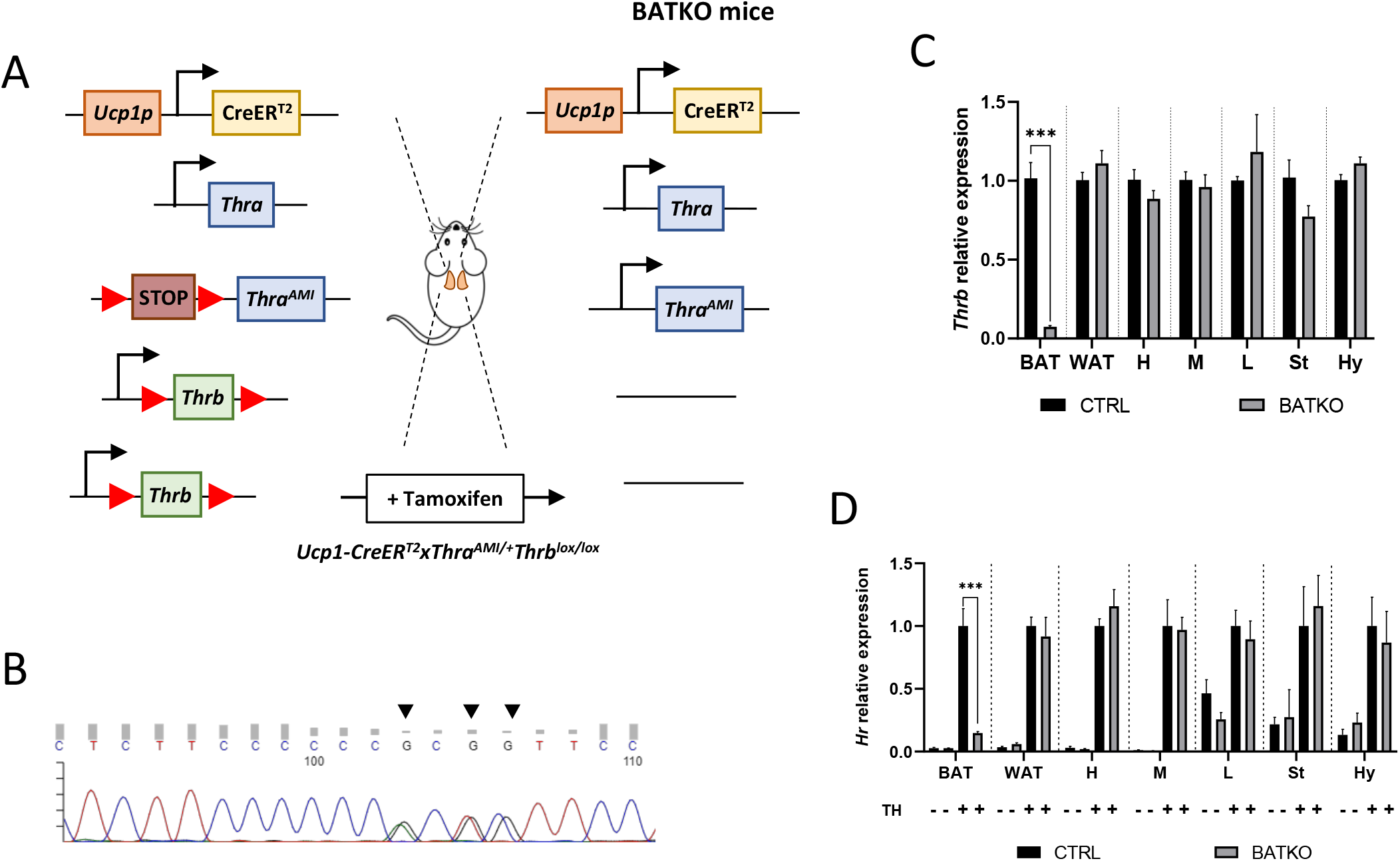
BATKO mice present a BAT-specific blockade of T3 signaling. **(A)** Schematic representation of the BATKO mice. BATKO mice carry the *Ucp1-CreER^T2^* transgene, allowing the brown adipocytes specific expression of the tamoxifensensitive CreER^T2^ recombinase. BATKO mice are also heterozygous for the *Thra^AMI^* allele, which encodes the TRα1^L400R^ dominant-negative receptor after Cre-mediated deletion of a STOP cassette flanked by loxP sequences. BATKO mice are homozygous for the *Thrb^lox^* allele in which exon 3 is flanked by two tandem-arranged loxP sequences. After tamoxifen injection, Cre-mediated recombination selectively excise the loxP-flanked sequences in brown adipocytes, resulting in the expression of TRα1^L400R^ and elimination of TRβ. Control mice (CTRL, not represented here) had the same genotype except for the absence of the *Ucp1-CreER^T2^* transgene and were also tamoxifen-treated. **(B)** Sanger sequencing chromatogram of a fragment of *Thra* cDNA prepared from BAT RNA of BATKO mice. Arrows indicate the positions of the TRα1^L400R^ mutations. **(C)** Relative mRNA expression of *Thrb* in different peripheral and central tissues of CTRL and BATKO mice (H: heart, M: muscle, L: liver, St: striatum, Hy: hypothalamus) (n=5-6/group). **(D)** Evaluation of T3 response in PTU-fed CTRL and BATKO mice through the induction of *Hr*, a well characterized TR target gene, after 24h of TH (+ or -; n=5-7/group). Statistical significance is shown for the comparison of CTRL and BATKO mice treated with TH. Error bars represent the SD. ***p < 0.001 for the indicated comparisons.

We verified that the *Thra^AMI^* allele was expressed in the BAT of BATKO mice (Figure 1B) and that *Thrb* expression was drastically reduced (93%) in the BAT of BATKO mice, but not in other tissues (Figure 1C). As expected, the T3-induced regulation of *Hr* expression, a classical T3 target gene in many tissues ^24,25^, was selectively and almost completely lost in the BAT of BATKO mice (Figure 1D). The observed residual responses most likely reflect the presence in BAT of other cell types, like endothelial cells and immune cells ^26^. Importantly, there was no significant difference between BATKO and CTRL mice in free T3 or free T4 serum levels, whatever the temperature and diet conditions used in this report (Suppl. figure 2). Altogether, BATKO mice present an altered T3 signaling specifically in brown adipocytes, allowing us to decipher the cell-autonomous functions of T3 in this cell type.

### Establishment of a catalog of TR direct target genes in BAT

We first aimed at establishing a complete list of TR direct target genes in brown adipocytes, defined as genes: 1) with a TR binding site (TRBS) within 30kb of the transcription start site (TSS) ^27^, 2) which mRNA levels are rapidly (within 24h) increased in BAT in response to T3 and T4 (collectively TH) and 3) which induction is lost in BATKO mice, *i.e*., controlled locally by TR in brown adipocytes (and thus, not secondary to the sympathetic stimulation of the tissue caused by the TH treatment). *Ucp1-CreER^T2^xThra^GS/+^mice* express specifically in brown adipocytes a GS-tagged version of TRα1 ^28,29^ used for ChIPseq. We identified 4210 TRBS, in the vicinity of 2311 genes, essentially located within 10kb of the TSS (Figure 2A) and mostly in intronic sequences (Figure 2B). TRBS were preferentially present on motifs related to the so-called DR4 consensus sequence (AGGTCAnnnnRGGnCA), described as preferential for TR fixation ^14^ (Figure 2C).

**Figure 2:**
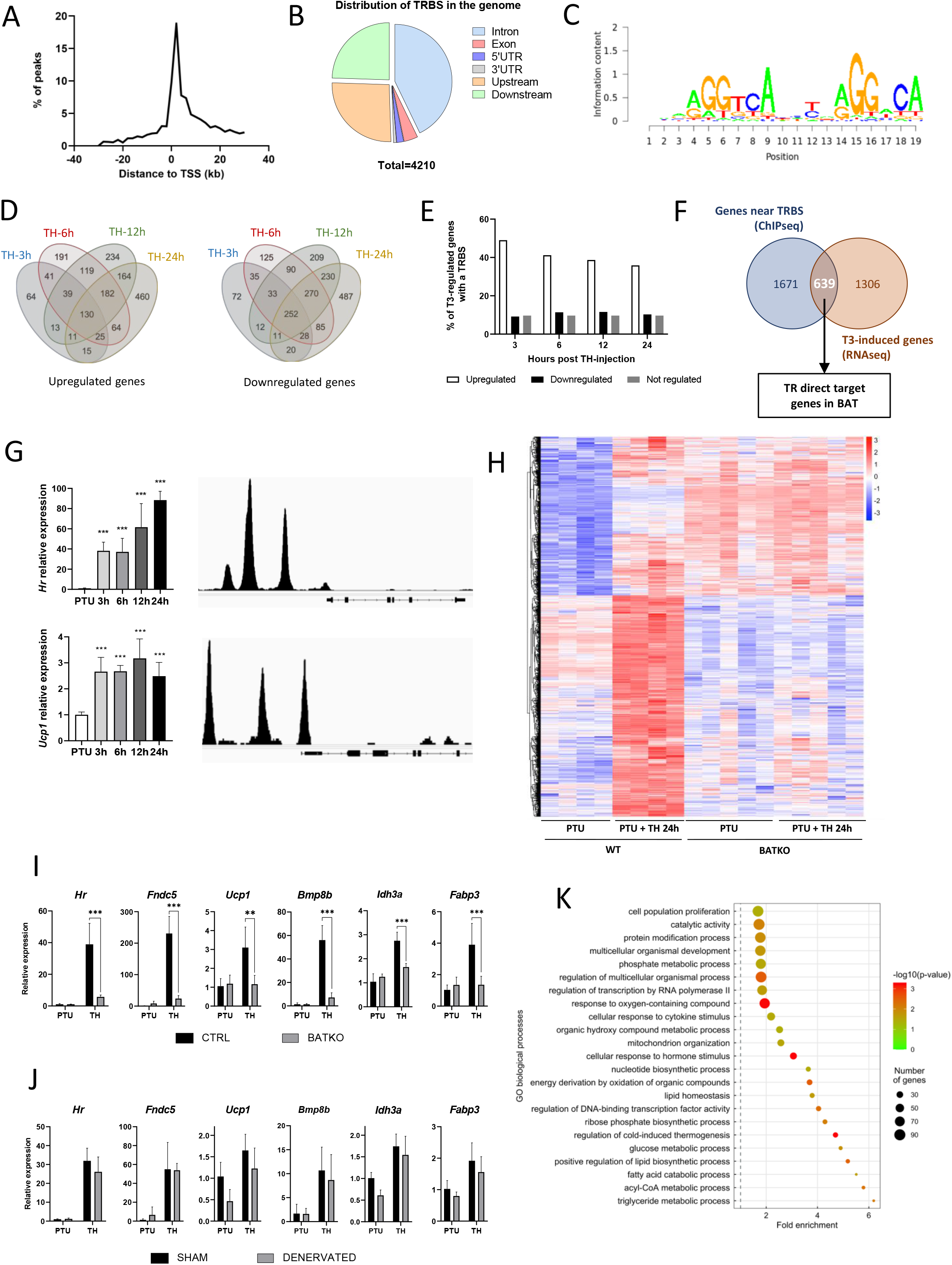
Identification of T3/TR target genes in brown adipocytes. **(A)** Consensus sequence found in BAT TRBS (Thyroid hormone Receptor Binding Site), as identified by *de novo* motif search. **(B)** Frequency of TRBS distribution around transcription start sites (TSS). **(C)** Pie chart of the 4210 TRBS distribution in the genome (UTR: untranslated region). **(D)** Venn diagrams of upregulated (left panel) and downregulated (right panel) genes after TH intraperitoneal injection in wildtype PTU-treated mice for different periods. **(E)** Percentage of genes which possess a TRBS within 30 kb of their TSS among genes whose expression in the BAT is regulated or not by T3 (upregulated in white, downregulated in black, not regulated in grey). **(F)** Venn diagram of genes whose expression is induced by T3 in at least one of the timepoints (in brown, RNAseq data) and genes with a TRBS within 30 kb of their TSS (in blue, ChIPseq data), i.e, TR direct targer genes. **(G)** Left: Time-course analysis of *Hr* (top) and *Ucp1* (bottom) expression in BAT after 24h of TH treatment of wild-type hypothyroid mice (RT-qPCR). Statistical significance is shown for the different time points *versus* untreated PTU-fed mice (n=4-6/group). Right: Extract of the TRBS in the *Mus musculus* genome browser around *Hr* (top) and *Ucp1* (bottom). **(H)** Heatmap representing in both CTRL and BATKO mice the expression of TR direct target genes up-regulated after 24h of TH injection in CTRL hypothyroid mice. Colors represent the *z-scores*, see scale besides the heatmap (n=4-5/group). **(I)** and **(J)** Relative expression of several TR direct target genes 24h after TH treatment in (I) CTRL/BATKO and (J) SHAM/DENERVATED PTU-fed C57BL6/J mice. Statistical significance is shown for the comparisons between CTRL-TH and BATKO-TH, or SHAM-TH and BATKO-TH mice (n=5-7/group). **(K)** Gene ontology dot plot of the 639 TR direct target genes. Only the ‘biological processes’ terms with a fold-enrichment > 1.5 were kept. Some of the terms were shortened to increase readability without affecting the meaning. Error bars represent the SD. **p < 0.01 and ***p < 0.001 for the indicated comparisons.

Time-course analysis of BAT transcriptome in hypothyroid (PTU-fed) wild-type mice treated with TH for 3, 6, 12 or 24h was conducted by RNAseq. It revealed that a large number of genes (1946 upregulated, 1744 downregulated) were regulated in a time-dependent manner (Figure 2D). We observed that 49% of genes induced by T3 after 3h possess a TRBS within 30kb of their TSS. This ratio fell below 10% for T3-repressed genes and for genes which expression was insensitive to T3 (Figure 2E). This suggests that downregulation of gene expression is not directly exerted by T3-bound TR, but is an indirect consequence of the TH treatment, either cell-autonomous or resulting from sympathetic stimulation. Given these considerations, the rest of the study was restricted to positively regulated genes.

We then crossed the RNAseq and the ChIPseq datasets to obtain a curated list of 639 putative TR direct target genes (Figure 2F, Data S1), whose expression was most likely under the direct control of T3/TR signaling. As expected, this set of genes included two emblematic TR target genes: *Ucp1* ^30^ and *Hr*^25^ (Figure 2G). Heatmap representation showed that TR direct target genes previously found to be upregulated after 24h of TH treatment in wild-type mice were not responsive in BATKO mice (Figure 2H). Some transcriptome results were confirmed by RT-qPCR for some of the identified TR direct targets (Figure 2I). We hypothesized that the residual response to T3 of BAT in BATKO mice might result either from the stimulation of brown adipocytes the sympathetic system ^21^ or from the T3 response of BAT cells which do not express *Ucp1*, like endothelial or immune cells. To distinguish between these two possibilities, we chemically destroyed BAT noradrenergic terminals of wild-type PTU-fed mice (Suppl. figure 3), treated or not afterwards with TH. T3-induced expression of target genes was mostly similar between sham and denervated mice (Figure 2J), indicating that, for the genes that we tested, the sympathetic stimulation was not involved.

According to gene ontology analysis, TR direct target genes were directly involved in the “regulation of cold-induced thermogenesis” (Figure 2K), including *Ucp1* and *Ppargc1a*, two fundamental actors of this process. Interestingly, *Ppargc1a* encodes for PGC1α, a co-activator of TR ^31^. Using a published list of PGC1α binding sites (GSE110053) ^32^, we found that around 33% of TR direct target genes showed a co-localization of TRBS and PGC1α binding sites (Suppl. figure 4), including genes involved in lipid metabolism as well as *Ucp1* and *Ucp3* (Data S2). In addition, many of TR direct target genes we identified were involved in mitochondrial transport, respiratory chain, catabolism and biogenesis of lipids, as well as genes involved in the glycolysis and the citric acid cycle. Finally, we also found genes involved in proliferation, a process of BAT mid-term adaptation to physiological stressors like cold ^6^. In summary, TR direct target genes belong to several biological processes, many of them being directly connected to BAT thermogenesis.

### Altered response of BATKO mice to temperatures below thermoneutrality

Based on the roles of TR target genes identified above, we predicted that BATKO mice would display alterations in BAT thermogenesis. At 23°C, neither body weight, nor body composition, nor metabolic rate were altered in BATKO mice, but food consumption was increased (Suppl. figure 5). As these small variations occurred at 23°C, which is a moderate cold exposure for mice ^33^, we submitted them to a more drastic cold challenge.

BATKO mice maintained their body temperature normally at 4°C during 72h (figure 3A) but they tended again to consume more food than CTRL mice (Figure 3B). We thus combined cold exposure with fasting causing a severe hypothermia in BATKO mice, which led us to end the experiment (Figure 3C). Thus, blockade of TH signaling in BAT prevents an adequate maintenance of body temperature in absence of a higher energy intake. This suggests that compensatory thermogenic processes are triggered to maintain body temperature in BATKO mice fed *ad libitum*. In that context, we notably observed that the browning of the WAT was exacerbated in BATKO mice (Suppl. figure 6).

**Figure 3:**
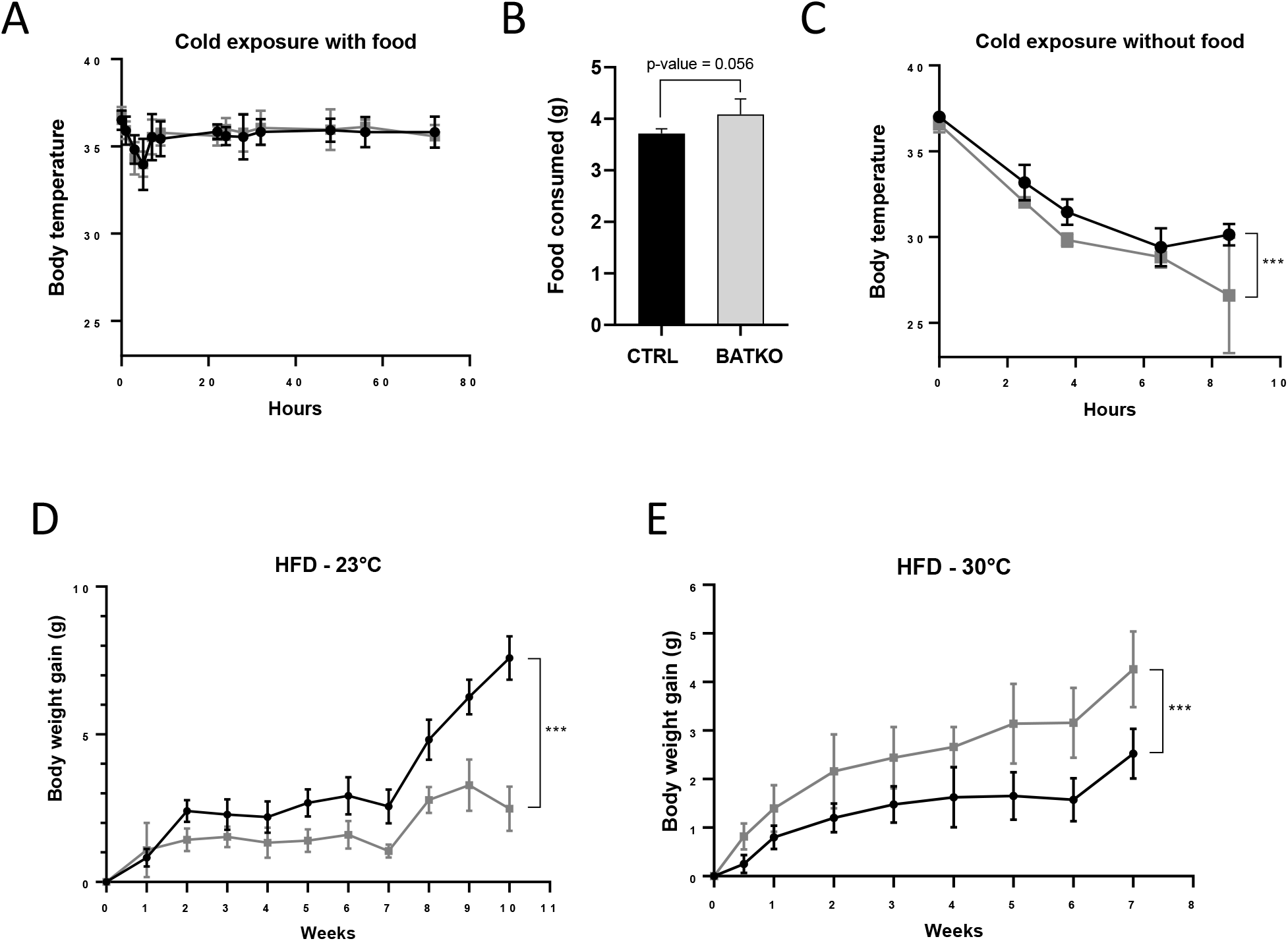
BATKO mice are cold-sensitive. **(A)** Core body temperature of CTRL and BATKO mice exposed to 4°C in presence of food (n=5-7/group). **(B)** Food consumption of CTRL (black) and BATKO (grey) mice exposed to 4°C (48h) (n=4/group). BATKO mice tended to eat more than CTRL mice. **(C)** Core body temperature in CTRL (black) and BATKO (grey) mice exposed to 4°C in the absence of food. After 8 hours of cold exposure, BATKO mice reached severe hypothermia and the experiment was stopped (n=4-5/group). **(D)** and **(E)** Body weight gain of CTRL (black) and BATKO (grey) mice at room temperature (D, n=4-5/group) or 30°C (E, n=5-8/group). Error bars represent the standard error of the mean. ***p < 0.001.

High-fat diet represents another challenge for the thermogenic capacity. BATKO mice proved to be resistant to diet-induced obesity at 23°C (Figure 3D) with similar food intake, suggesting higher energy expenditure. Higher energy expenditure was not observed by 48h of indirect calorimetry (Suppl. figure 5) but even a minor difference could explain the subtle difference between BATKO and CTRL mice. HFD feeding was then conducted at thermoneutrality, eliminating the need to activate alternate mechanisms to defend body temperature. In this condition, the opposite result was observed, BATKO mice being more sensitive to diet-induced obesity than CTRL mice (Figure 3E). Collectively, these results point out that BATKO mice suffer from a reduced efficiency of BAT adaptive thermogenesis both in condition of cold exposure or excess of calories. When exposed to HFD at 23°C, the activation of alternative thermogenic processes combined with the defect in adipocytes thermogenesis results in a paradoxical resistance to obesity of BATKO mice.

### BAT TR-signaling controls the expression of a subset of genes induced during cold exposure

To better understand the molecular response controlled by TR signaling in the BAT during cold response, we compared the BAT transcriptome of BATKO and CTRL mice after 24h at 4°C, in the presence of food. Among 2865 cold-induced genes (Data S3), 491 (17%) displayed a different response in BATKO mice (Figure 4A). For most of them, the cold induction was partially or completely lost in BATKO mice. Noteworthy, *Ppargc1a* was the only classical thermogenic marker that was part of this subset (Data S3). The highly stringent statistical interaction model that we used ^34^ failed to reveal an influence of T3 on the cold-response of the *Ucp1* and *Dio2* genes, two classical thermogenic markers. Thus, we used RT-qPCR to measure *Ucp1* and *Dio2* mRNA levels on a larger number of samples and found that the cold-induction of these genes was indeed altered in BATKO mice (Figure 4B). Thus, the high stringency of the statistical model used avoids false positives, allowing to trustfully highlights genes of interest, but some of them can be missed due to a lack of statistical sensitivity.

**Figure 4:**
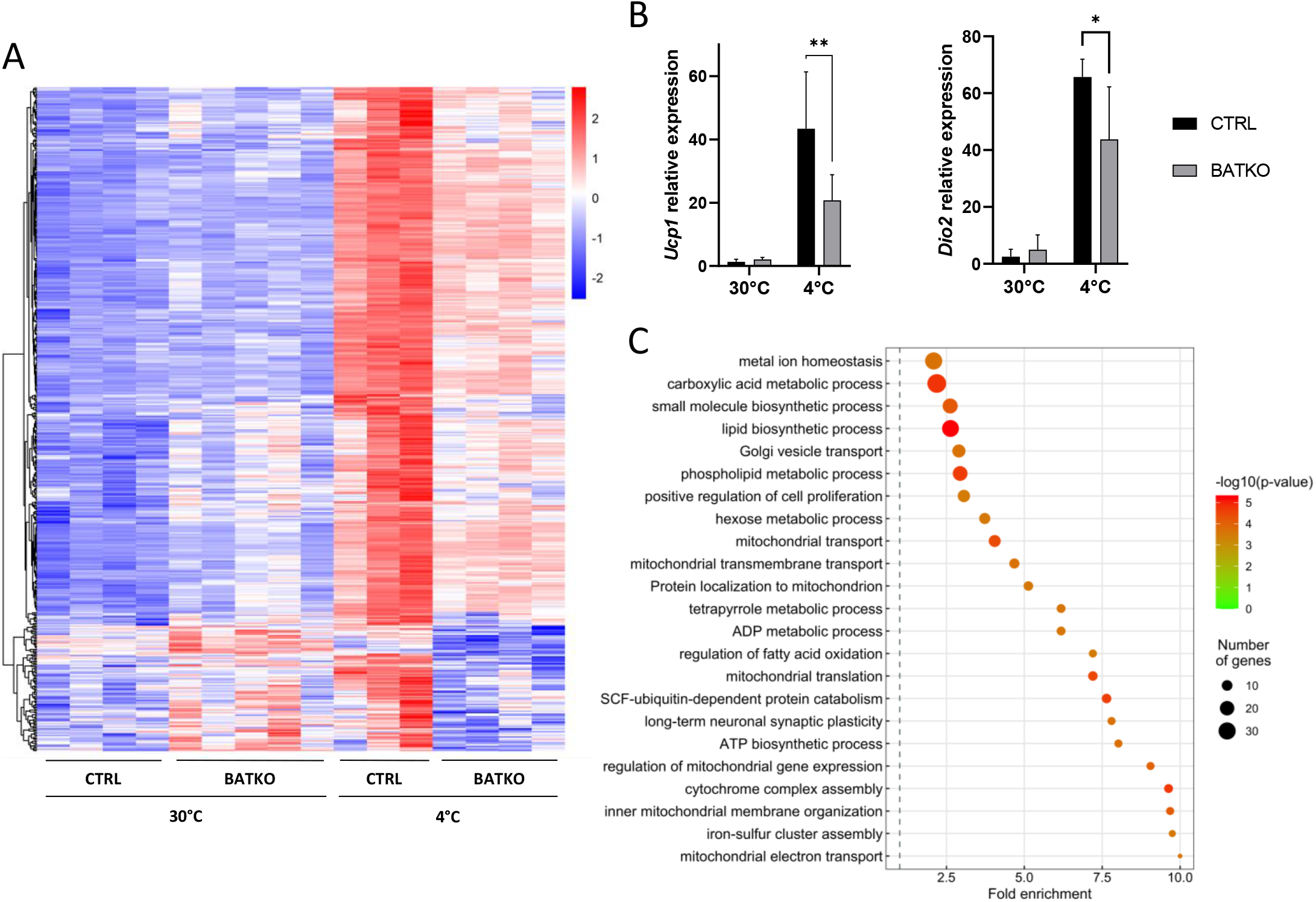
A subset of T3-regulated genes is activated during cold exposure and is necessary for an efficient BAT thermogenic response. **(A)** Heatmap representation of cold-responsive genes altered in BATKO mice, after 24h at 4°C. Colors represent the z-scores, see scale in heatmap (n=3-5/group). **(B)** Relative mRNA expression of *Ucp1* (left panel) and *Dio2* (right panel) in CTRL and BATKO mice at 30°C or after 24h at 4°C. Statistical significance is shown for the comparison between CTRL and BATKO mice at 4°C (n = 4-6/group). **(C)** Gene ontology dot plot representation of biological processes enriched in the 491 genes inefficiently induced in BATKO mice at cold. Some of the terms were shortened to increase readability without affecting the meaning. Error bars represent the SD. *p < 0.05, **p < 0.01 for the indicated comparisons.

Using gene ontology, we found that the genes which cold-response was significantly altered in BATKO mice were often directly connected to thermogenesis (Figure 4C). This gene set notably included genes involved in mitochondrial activity and respiratory chain. Other genes were involved in glycolysis, Krebs cycle, lactate metabolism and glucose transport which have all been shown to be crucial to fuel BAT thermogenesis ^35^. Genes involved in both lipolysis/fatty acid oxidation and lipogenesis, two processes required for an appropriate lipid use during thermogenesis, were also altered. Finally, a gene set reflecting cell proliferation was activated (Data S4). Restriction of this list to the TR direct target genes (Suppl. figure 7) also highlighted secreted peptides (*Bmp8b, Fgf1*) and enzymes participating in heat-producing creatine futile cycle (*Alpl). Ppargc1a* still belonged to this list of restricted genes of interest, reinforcing its importance in TR-mediated regulation, as abovementioned. Collectively, the BAT transcriptome of BATKO mice evidenced several altered pathways that are crucial for BAT thermogenesis, which could explain their phenotype.

### T3 in BAT controls BAT proliferation

RNAseq analysis revealed that TH might participate in BAT proliferation, both during hyperthyroidism and cold exposure. After 5 days of TH treatment, wild-type PTU-fed mice displayed an overexpression of 72% of a group of genes previously described to be involved in cell-cycle ^36^ (Suppl. figure 8). This translated into an effective increase in cell proliferation when TH treatment was associated with an injection of EdU 24h before sampling. This effect was insensitive to BAT denervation (Suppl. figure 9), suggesting that it most likely results from a cell-autonomous response to T3.

As predicted by the transcriptomic data after 24h of cold exposure, we also observed that proliferation is triggered in the BAT when CTRL mice were exposed 72h to cold, but 2.5-fold less proliferative cells were observed in BATKO mice, in the same conditions (Figure 5).

**Figure 5:**
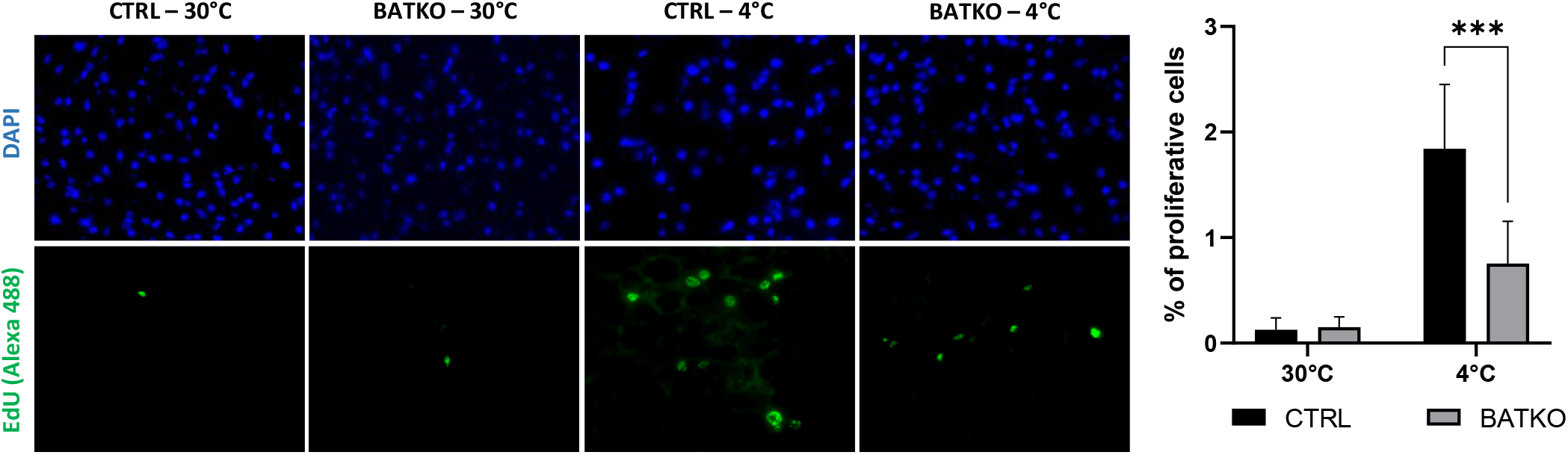
Local control of brown adipocytes proliferation by T3. Representative images (left) and quantification (right) of EdU-positive proliferative cells in BAT from both CTRL and BATKO mice exposed 72h to 4°C and injected with EdU after 24h and 48h of cold. Nuclei are stained in blue with DAPI. Percentage of proliferative cells is the ratio of proliferative cells on the number of nuclei (n=4-7/group). Statistical significance is shown for the comparison CTRL 4°C *versus* BATKO 4°C. Error bars represent the SD. *p < 0.05 and **p < 0.01 for the indicated comparisons.

As most of the cell-cycle genes induced by T3 do not have a TRBS within 30kb (Suppl. figure 8A), the link between the TR direct target genes and proliferation in BAT yet remains uncertain. *Ccnd1*, encoding cyclin D1, is one of the TR direct target gene (Data S1) and also belongs to the genes that are under the control of TH signaling during cold exposure. It is thus an interesting candidate to play a significant part in this process.

## Discussion

T3 has been for a long time under the lights of metabolism research for its ability to regulate energy expenditure in many tissues, including the BAT. However, neither TR direct target genes in BAT, nor the contribution of BAT in T3-mediated regulation of energy expenditure have been elucidated so far. Here, we present an unprecedented in-depth analysis of T3 direct influence on gene expression in brown adipocytes and developed a transgenic mice model with a brown-adipocyte specific suppression of T3 signaling. To our knowledge, this model is currently the only one allowing a suppression of T3 signaling specifically in brown adipocytes at adult stages. This prevents any BAT development alteration, a limitation met in other non-conditional mice models ^37^. Collectively, it showed that T3 signaling was pivotal for BAT adaptive thermogenesis, regulating the management of thermogenic fuels as well as the plasticity of the tissue.

Combined with transcriptome analyses, the genome-wide study of TRα1 chromatin binding allowed us to identify 639 genes whose transcription is most likely controlled by liganded TR in brown adipocytes. This set of genes only partially overlaps with the ones described in other cell types, probably as a result of differential chromatin occupancy ^27–29^ and the presence of cell-specific transcription cofactors. One of these cell-specific cofactors is PGC1α (encoded by *Ppargc1a*) which we found here to be a TR direct target gene. As PGC1α is a transcriptional coactivator of TR ^31^, it confirms previous hypothesis that TR and PGC1α are involved in an auto-regulatory feed-forward loop ^38^. In line with this, we found that 33% of the TRBS in brown adipocytes are also occupied by PGC1α, notably for genes involved in lipid metabolism or directly involved in thermogenesis, like *Ucp1* and *Ucp3* ^39,40^. PGC1α is thus likely to play a pivotal role in the T3-dependent regulation of energy metabolism in brown adipocytes. As PGC1α is also a coactivator of several other nuclear receptors, notably PPARγ and ERRα, its overexpression might generate a cross-talk with other signaling pathways.

One of the main challenges of our study was to make a distinction between a local influence of T3 and an indirect consequence, resulting from the hypothalamus response to T3 ^21^. The BAT response to T3 is largely lost in BATKO mice and conserved in denervated mice, suggesting that the changes in gene expression observed in BAT mainly reflect the cell-autonomous response initiated by TR in brown adipocytes and not the effect of T3 mediated by the hypothalamus ^21^. Residual responses to T3 in BATKO may reflect the sensitivity of other cells types present in the BAT which do not express the *Ucp1-CreER^T2^* transgene or an incomplete effect mediated by tamoxifen. Otherwise, they could be secondary to a modification of circulating peptides caused by the T3 treatment on other organs. For instance, *Fgf21* expression is induced by T3 in the liver ^41^ and the resulting secreted peptide has been shown to promote BAT thermogenesis ^42^. We found indication for a reciprocal influence as several TR direct target genes in brown adipocytes encode secreted factors and adipokines (*Apln, Fdcn5, Bmp8b*), that are able to act in other organs. This suggests that T3 can influence a complex network of cross-talks in the organism.

T3 signaling in BAT is triggered in CTRL mice during cold exposure as it was previously shown by the activation of type 2 deiodinase activity in similar conditions ^43^. This activation of T3 signaling represents a significant fraction of the cold response, since 17% of the gene regulations induced in the BAT by cold exposure in CTRL mice are lost in BATKO mice. We showed that T3 signaling in the BAT controls the expression of genes crucial for both lactate metabolism (*Ldha*) or glycolysis (*Hk1, Aldoa, Pkm, Pfkl*), two pathways essential for BAT thermogenesis ^35^. T3 also controls genes involved in lipid metabolism, which is also congruent with previous observations revealing that T3 signaling is required for efficient lipogenesis ^44^ and lipolysis ^45^. Here, we bring a broader picture by identifying all the genes that are regulated by T3 for these processes.

Interestingly, we showed that *Ppargc1a* was among impacted genes. As it coactivates other nuclear receptors, we can imagine that part of the genes dysregulated in BATKO mice can be attributed to other signaling pathways downstream of PGC1α. On the other hand, we observed that other classical thermogenic markers were not the most obviously affected in BATKO mice. Indeed, *Ucp1* and *Dio2* could not be detected as differentially expressed by our stringent whole-transcriptome approach but only by targeted expression assessment. Among TR-dependent cold-induced genes, we rather detected *Alpl*, a phosphatase required in brown adipocytes for UCP1-independent thermogenesis derived from futile creatine cycle ^46^. Thus, T3 signaling in BAT coordinates the expression of genes involved in both UCP1-dependent and UCP1-independent thermogenesis. Finally, we showed that T3 signaling in brown adipocytes controls cell proliferation both during cold exposure or upon hyperthyroidism. This concurs with a recent study showing that T3 promotes BAT proliferation in adult mice ^47^. Also, the link between TR target genes and the cell cycle remains uncertain. The direct activation of *Ccnd1*, encoding cyclin D1, provides a hypothetical link between T3 and proliferation. This intertwining might also occurs through the regulation of cell cycle gene regulators, as already demonstrated for CREB and AP-1 ^48^.

Thus, inefficient handling of metabolic fuels, reduction of the tissue plasticity and defect in thermogenic biochemical pathways can explain the apparent default in BAT thermogenesis. This can be compensated by an increased energy consumption which could explain why BATKO mice gained less weight than CTRL mice at room temperature, which represents a mild cold challenge ^49^. This extra-energy consumption might favor the onset of other thermogenic mechanisms, as observed with the exacerbated WAT browning. However, WAT browning’s ability to consume a significant amount of energy has recently been called into question ^50^. Although we failed to measure a striking difference in BATKO mouse oxygen consumption at room temperature over a 48h-period, a minor increase in the respiratory quotient might have long-term consequences. At thermoneutrality, where no compensatory mechanisms are required to maintain body temperature, BATKO mice became hypersensitive to diet-induced obesity. This reflected the alteration of diet-induced BAT thermogenesis and the inability of this tissue to optimally use metabolic substrates and thus expend energy. A similar phenotype was previously observed for other mouse models deficient for BAT thermogenesis ^51^.

In conclusion, we used a combination of both omics data and BAT-specific TR-signaling alteration in mice to introduce an unprecedented view of T3 cell-autonomous response in brown adipocytes. We showed that T3-signaling in BAT controls both key metabolic pathways and tissue plasticity that are essential for adaptive thermogenesis. These results represent a valuable database that pave the way for further metabolic studies to detail the molecular implications of T3 signaling during BAT adaptive thermogenesis. Finally, as T3 acts in many other tissues, a similar approach could be used to put together the puzzle of T3 influence in energy expenditure.

### Limitations of the study

One of the objectives of this study was to define a catalog of TR direct target genes in brown adipocytes, based on a combination of RNAseq and ChIPseq. Both TRα and TRβ are present in the BAT ^13^ but we only performed the TRα1 ChIPseq due to the absence of adequate tools for the TRβ ChIPseq. The cistromes of the two receptors might not fully overlap, and we might have missed a fraction of the TRBS on the genes of interest. However, this limitation should be tempered since previous data have indicated that TRβ-selective binding site are infrequent in the mouse genome ^27^. Another limitation is that we assigned genes to a given TRBS by considering only its distance to the transcription starting site. Chromatin 3D organization and insulation are well known to influence the interactions between distant regulatory elements, and the linear distance is only partially informative. The additional genomic investigations required to overcome these limitations remain for the moment hardly feasible in mouse tissues.

## Materials and Methods

### Animal procedures

All experiments were carried out in accordance with the European Community Council Directive of September 22, 2010 (2010/63/EU) regarding the protection of animals used for experimental and other scientific purposes. The research project was approved by a local animal care and use committee (C2EA015) and authorized by the French Ministry of Research.

### Genetically modified mouse models

The genetic background of all mice that were used in the present study was C57BL6/J. *Thra^AMI/+ 22^, Thrb^lox/lox^* ^23^ and *Ucp1CreER^T2^* ^52^ mouse lines were crossbred to introduce the different recombinant alleles in *Ucp1CreER^T2^xThra^AMI/+^Thrb^lox/lox^* mice. The expression of the *ThraA^MI^* allele allows the expression of the TRα1^L400R^ mutant, which has dominant-negative properties over TRα and TRβ, after Cre/loxP-mediated excision of a stop cassette ^22^. Despite the persistence of one intact *Thra* allele in *Thra*^AMI/+^ mice after Cre-recombinase action, the dominant-negative action of TRα1^L400R^ eliminates the capacity of cells to respond to T3. *Thrb^lox^* has 2 tandem-arranged loxP sequences, allowing Cre-mediated excision of exon 3 of the *Thrb* gene, which encodes the DNA binding domain of the TRβ1/TRβ2 receptor, resulting in a frameshift and a loss of function ^23^. In the present study, we used mice of the *UCP1CreER^T2^* line, in which CreER^T2^ expression is under the control of the *Ucp1* promoter ^52^. In the absence of physiological stressors, *Ucp1* is specifically expressed in brown adipocytes, thus restricting Cre-mediated recombination to this cell type. CreER^T2^-recombinase action occurs after its translocation to the nucleus, allowed by tamoxifen treatment. Tamoxifen was injected intraperitoneally every day during 5 days at 50 mg/kg of mice.

In *Ucp1CreER^T2^* mice, we used a *ROSA26tdTomato* reporter transgene, also known as *Ai9* (Suppl. figure 1) ^53^, we verified the absence of recombination activity outside BAT, except for a small fraction of cells present in the choroid plexus. In the following, mice with the *Ucp1CreER^T2^xThra^AMI/+^Thrb^lox/lox^* genotype were called BATKO. *Thra^AMI/+^Thrb^lox/lox^* littermates, also injected with tamoxifen, were used as controls (CTRL).

In order to address chromatin occupancy by TR specifically in brown adipocytes, we used *Thra^GS/+^* mice ^28^ to generate *ad hoc Ucp1CreER^T2^xThra^GS/+^* transgenic mice. These mice were generated by knocking in the *Thra* locus a sequence encoding TRα1 fused with protein G and streptavidin protein (GS) after a floxed stop cassette. In the presence of the *Ucp1CreER^T2^*, tamoxifen injection allows the recombination at the loxP sites that excise the stop cassette specifically in brown adipocytes, allowing the expression of the GS-TRα1 only in this cell-type. This strategy has already been successfully used in other tissues, including striatum and heart ^29,28^.

### Experimental animal procedures

We used 2-to-5-month-old male mice for experiments. Genetically modified mice were generated in our own animal facility, whereas wild-type C57BL6/J mice were ordered from a commercial supplier (Charles River). Mice were fed *ad libitum* with LASQC Rod16 R diet (Altromin, Germany) and housed under recommended conditions (notably, at room temperature, i.e., 23°C). Hypothyroidism in adult animals was induced as previously described, with 14 days of treatment with a propylthiouracil (PTU)-containing diet (Harlan Teklad TD95125, Madison, WI) ^54^. It was combined in some cases by hyperthyroidism induced by intraperitoneal injections of a T3/T4 mix (T4 at 2*μ*g/g of mice and T3 at 0.2 *μ*g/g of mice, Sigma-Aldrich), daily for the five last days or once at day fourteen of the PTU treatment. T3, the active compound, was not injected alone to get close to hyperthyroid conditions met *in vivo*.

Before assessment of cold response, mice were housed for 10 days at 30°C (with normal 12h light/dark cycles) and subcutaneously implanted with IPTT-300 transponders (Plexx BV, Netherlands). The mice were then housed in pairs without enrichment and placed at 4°C during the indicated period. Infrared thermography was performed on awake mice after 48h of cold exposure, using an infrared camera (FLiR Systems, Inc). For cell proliferation assessment, EdU (Thermofisher) was intraperitoneally injected twice (100mg/kg, after 24h and 48h of cold exposure) and mice were killed after 72h of cold exposure. This allowed to stain proliferative cells over the last 48h of cold exposure.

At the end of experiments, mice were anesthetized by an intraperitoneal injection of xylazine (25mg/kg) and ketamine (130mg/kg) mixture. Blood was drawn from the vena cava and collected in heparin-coated tubes to retrieve plasma. Several tissues and organs were dissected and either directly processed for histology or snap frozen and stored at −80°C for later RNA preparation.

### Indirect calorimetry and body composition measurement

Body composition was measured in awake mice by low-field nuclear magnetic resonance with a Minispec LF90II device (Bruker). Phenomaster metabolic cages were used for indirect calorimetry measurements (TSE Systems, Berlin, Germany). Mice were first placed in individual cages for a 24h period of habituation to isolation, after which oxygen consumption (VO_2_) and carbon dioxide rejection (VCO_2_) were continuously recorded for 48h, under a normal 12h light/dark cycle. VO_2_ and VCO_2_ were expressed as mL/hours/kg of mice. Respiratory quotient was obtained as the VO_2_ / VCO_2_ ratio. The metabolic rate was calculated according to the Weir formula ^55^ as following: Metabolic rate (kcal/min) = 3.94*VO_2_ + 1.1*VCO_2_.

### BAT denervation

Chemical denervation of BAT sympathetic nerve endings was performed in wild-type PTU-fed mice under isoflurane 2% anesthesia and ketoprofen (1mg/kg) analgesia. This technique, which involves injections of 6-hydroxydopamine (6-OHDA), a neurotoxin selective for sympathetic neurons, permits a specific denervation of sympathetic neurons, while keeping the sensory fibers intact ^56^. The effects of sympathetic denervation were compared to those obtained after vehicle injection (0.15mol/L NaCl and 1% ascorbic acid, sham mice). Briefly, the two lobes of interscapular BAT were exposed through a midline skin incision along the upper dorsal surface and gently separated from the skin with surgical forceps. Then, injections of 6-OHDA (Sigma-Aldrich) were performed directly into each lobe of the interscapular BAT. For each lobe, 10μL (10mg/mL) was injected in several times (10 injections of 1μL) using a Hamilton syringe (i.e., 20μL/mice). The skin incision was then closed with several surgical stitches. Animals were allowed to recover for 5 days before further experiments.

### Western-blot analysis

Cell extracts from BAT were lysed in standard lysis buffer (20mM Tris-HCl, pH 8, 138mM NaCl, 1% NP40, 2.7mM KCl, 1mM MgCl2, 5% glycerol, 5mM EDTA, 1mM Na3VO4, 20mM NaF, 1mM DTT, 1% protease inhibitors), and homogenized using FastPrep^®^ (MP Biomedicals). Proteins were assayed in triplicate with Pierce™ BCA Protein Assay Kit (Thermo Fisher Scientific). Aliquots of 30 μg of proteins, denatured in buffer (20% glycerol, 10% β-mercaptoethanol, 10% SDS, 62.5mM Tris) were analyzed from 12% SDS polyacrylamide gel electrophoresis and transferred to PVDF Immobilon membranes (Biorad). After 1h-saturation in TBS/0.2% Tween/2% milk at room temperature, the membranes were probed (overnight at 4°C) with rabbit polyclonal anti-tyrosine hydroxylase (Merck, AB152) diluted in TBS/0.2% Tween/2% milk. Then, membranes were rinsed threetimes in TBS/0.2% Tween for 10min and incubated for 1h with goat secondary anti-rabbit IgG linked to peroxidase (dilution 1:5,000; Biorad) in TBS/0.2% Tween/2% milk. Membranes were rinsed again and exposed to Clarity™ Western ECL Substrate (Biorad). The intensity of the spots was determined by densitometry with ChemiDoc Software (Biorad) and analyzed using the Image LabTM software (Biorad). Quantification of whole protein levels (using the stain-free protocol provided by Biorad) was used for normalization. Stain-Free technology enabled fluorescent visualization of 1-D SDS PAGE gels and corresponding blots. The relative amount of total protein in each lane on the blot was calculated and used for quantitation normalization.

### Histology

BAT and WAT samples were fixed in 10mL Zinc Formal Fixx (ThermoFisher Scientific) during 24h at 4°C. Samples were dehydrated using the Leica TP1020 semi-enclosed processor and embedded in paraffin. 6-μm sections were processed for immunohistochemistry (IHC) and EdU detection. Tissue sections from different mice were assayed on the same slide to minimize staining variability.

For IHC, deparaffinized sections were incubated overnight at 4°C with rabbit poly-clonal antibodies directed against UCP1 (Abcam, ab10983), diluted 1:400 in PBS/2.5% goat serum. Sections were then incubated with a horseradish peroxidase-labeled anti-rabbit antibody (1:300, Promega, W401B) for 1 hour at room temperature. Peroxidase activity was visualized with diaminobenzidine staining (DAB, Sigma-Aldrich, D5905). Images were acquired using an AxioObserver Zeiss microscope at a 16x magnification.

EdU detection was assessed as recommended by the manufacturer (Click-iT EdU Cell Proliferation Kit, Thermofisher), following deparaffinization of BAT sections. Images were acquired on a DM6000 Leica microscope. Both EdU-positive cells and DAPI-marked nuclei were quantified on pictures of whole BAT sections to avoid any bias of field selection.

### Plasma T3/T4 quantification

Plasmatic free T3 and free T4 were quantified on a Cobas 6000 automat with the Cobas e601module (Roche, ECL analysers).

### RNA extraction and RT-qPCR

RNAs were extracted using Trizol (Invitrogen, Carlsbad, CA, USA). Total RNA was reverse transcribed to cDNA using MMLV reverse transcriptase (Promega, Wisconsin, USA). RT-qPCRs were performed using SYBRGreen mix (BioRad iQ supermix). The results were analyzed according to the ΔΔCT method ^57^. *Hprt* was used as the reference gene. Primers are listed in Table 1.

**Table 1.**
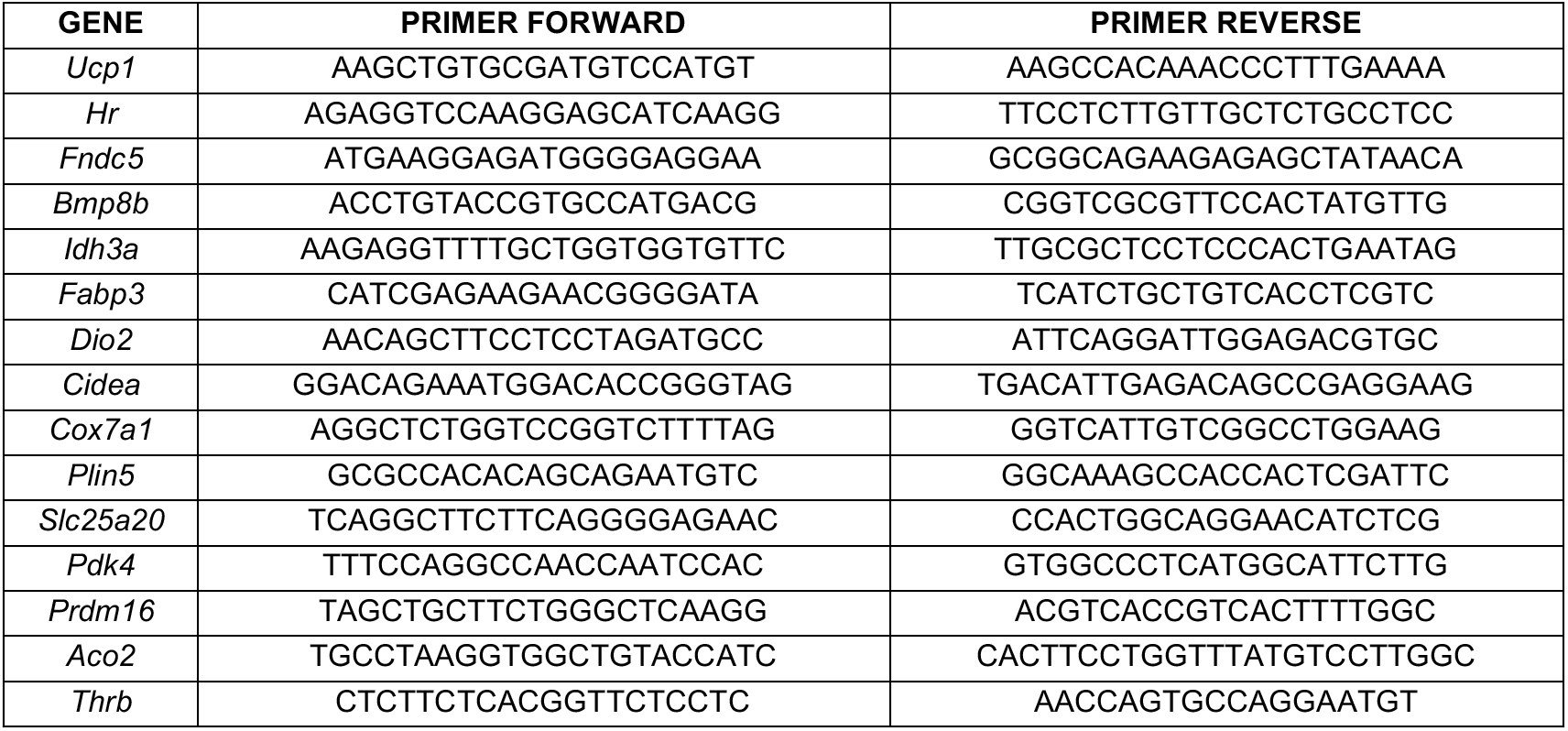
Primer sequences

### RNAseq analysis

cDNA libraries were prepared using the total RNA SENSE kit (Lexogen, Vienna Austria) and analyzed on a Nextseq 500 sequencer (Illumina) as previously described ^29,58^. Raw data of single-end sequencing were aligned on the GRCm38 (mm10) reference genome using Bowtie (Galaxy Version 2.2.6.2) and converted to count tables using htseqcount (Galaxy Version 0.6.1galaxy3) respectively. Differential gene expression analysis was performed with DESeq2 (R package, Version 1.34.0) ^34^ using the following thresholds: adjusted p-value <0.05; average expression > 10 reads per million, log_2_ fold-change > 0.6 or < −0.6. The effects of mutations on cold exposure were assessed using the interaction model ^59^. Thresholds for the interaction model were the following: average expression > 10 reads per million, adjusted p-value < 0.05.

Differentially expressed genes expression was visualized as clustered heatmaps using Pheatmap R package (RRID:SCR_016418). Normalized counts from DESeq2 were used as inputs and the correlation method was used to cluster the genes. Data were scaled independently for each gene, with the same color code for all genes (red: above mean, white: mean, blue: below mean).

Gene ontology analyses were made using the Gene Ontology Resource (http://geneontology.org).

### Chromatin immunoprecipitation sequencing (ChIPseq) and analysis

Freshly dissected small pieces of BAT were incubated during 25min at room temperature under agitation with a crosslink solution (250mM disuccinimidyl glutarate, 50mM HEPES, 100mM NaCl, 1mM EDTA, 0.5mM EGTA). Then, 1% formaldehyde was added for 20 min followed by 50mM glycine addition. The tissue was rinsed several times with cold PBS. Chromatin immunoprecipitation was then performed as previously described ^27^. Sequencing libraries were prepared from the immunoprecipitated fraction and the input fraction as a control, using the Accel-NGS 2S Plus DNA library kits with single indexing (Swift Biosciences). They were analyzed on a Nextseq 500 sequencer (Illumina). Raw data of paired-end sequencing were aligned on the GRCm38 (mm10) reference genome using Bowtie (Galaxy Version 2.2.6.2). MACS2 (Galaxy Version 2.1.1.20160309.0) was used for peak calling and peaks with a score inferior to 60 were filtered out. *De novo* motif search was performed using SeqPos motif tool (version 1.0.0). Genes within 30kb of peaks were called out using GREAT (http://great.stanford.edu/public/html/). We chose a distance of 30kb upstream or downstream of the transcription start site (TSS) to attribute a TR binding site (TRBS) to a gene. Although arbitrary, this distance was found to maximize the ratio of T3-responsive genes among the included genes, without excluding genes which have been well characterized as TRα1 target genes in other neural systems, such as *Klf9* or *Hr* ^60^. The distribution of distances of TRBSs around TSSs, as well as the distribution of TRBSs in the genome, were assessed using PAVIS (https://manticore.niehs.nih.gov/pavis2/).

### Statistical analysis

The data shown represent the average values for animals with the same genotype that were given the same treatment. The number of animals used in each experiment (*n*) is indicated in figure legends. Except when anything else is mentioned, the error bars represent the standard deviation (SD). For comparing two means, the statistical relevance was determined using an unpaired Student’s t-test. For determining the effects of the mutations on food consumption, body temperature, body weight overtime, we used a two-way ANOVA with time as one factor, and the other parameter as the second factor. For comparing the different levels of one factor to a control group, the statistical relevance was determined using the one-way ANOVA method. When this test showed significant differences (*p*-value < 0.05), a *post-hoc* Tukey test was used for multiple comparisons. To assess the effects of the mutations or the denervation on a given treatment (2 factors with 2 levels each), the statistical relevance was determined using a two-way ANOVA. Only when the interaction term was significant (interaction *p*-value < 0.05), a *post-hoc* Tukey test was used to compare the effects of the indicated combinations. For any of these tests, statistical relevance is shown in the graphs as follows: * *p*-value < 0.05, ** *p*-value < 0.01, *** *p*-value < 0.001.

## Data availability

The raw sequencing data and aligned read counts generated as part of this study has been deposited to the NCBI Sequence Read Archive. Accession number: GSE201136; https://www.ncbi.nlm.nih.gov/geo/query/acc.cgi?acc=GSE201136.

## Acknowledgments

We thank Catherine Cerrutti and Gerard Benoit for the advices about sequencing analysis as well as Benjamin Gillet, Sandrine Hughes and the PSI platform of the IGFL for deep sequencing. We also thank Emmanuel Quemener from Center Blaise Pascal/ENSL for the development and maintenance of the ENS Galaxy portal with the help of SIDUS (Single Instance Distributing Universal System). We acknowledge the contribution of the SFR Santé Lyon-Est (UCBL, UAR3453/CNRS, US7/Inserm) facility ANIPHY for their help and the contributions of the CELPHEDIA Infrastructure, especially the center AniRA in Lyon and the Equipex, ANR-11-EQPX-0035 PHENOCAN. We thank Nadine Aguilera and the ANIRA-PBES facility for the mice transgenesis and breeding. We thank internship students Eleonora and Mathilde for their implication at the early steps of the project. We also thank Marine and Manon for their involvement in denervation experiments. Finally, we thank C. Wolfrum for providing *Ucp1*-CreER^T2^ mice. This work was supported by the European Union’s Horizon 2020 research and innovation program under grant agreement no. 825753 (ERGO).

---

**Supplementary figure 1:**
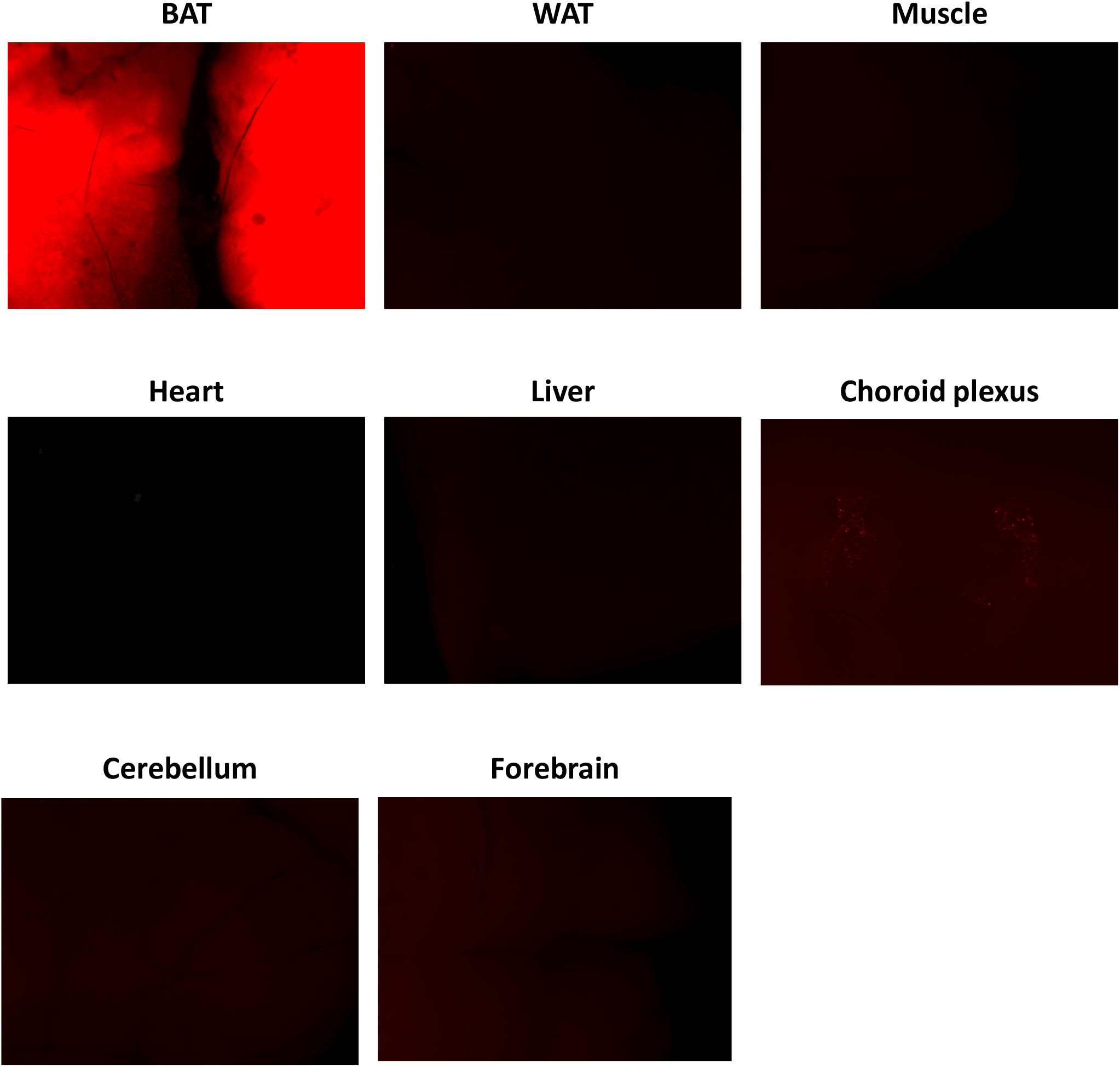
CRE activity in peripheral and central tissues. *Ucp1CreER^T2^:ROSATomato^lox^* reporter mice were dissected and their tissues observed under a Leica M205FA fluorescent stereomicroscope. Results are shown for one mouse.

**Supplementary figure 2:**
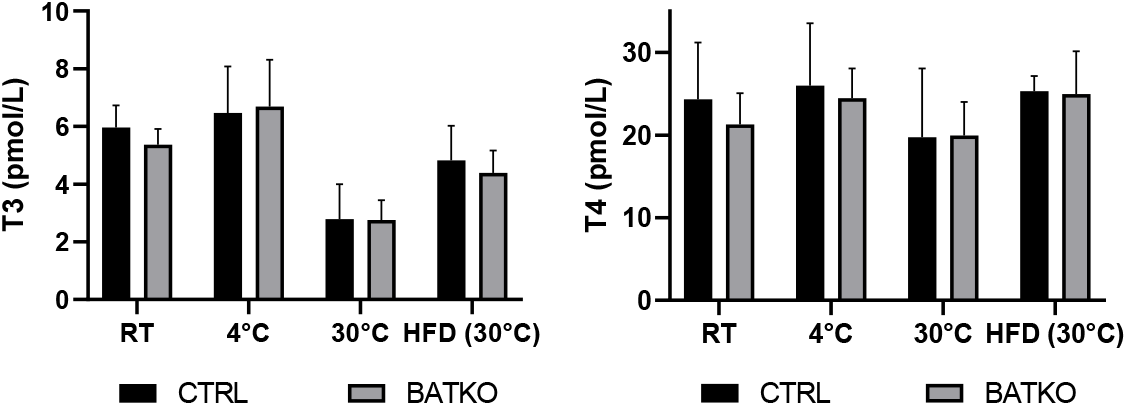
Serum concentration of TH in CTRL and BATKO mice. Serum concentration (in pmol/L) of free T3 (left panel) and free T4 (right panel) in CTRL and BATKO mice in different experimental conditions (RT: room temperature) (n=4-5/group).

**Supplementary figure 3:**
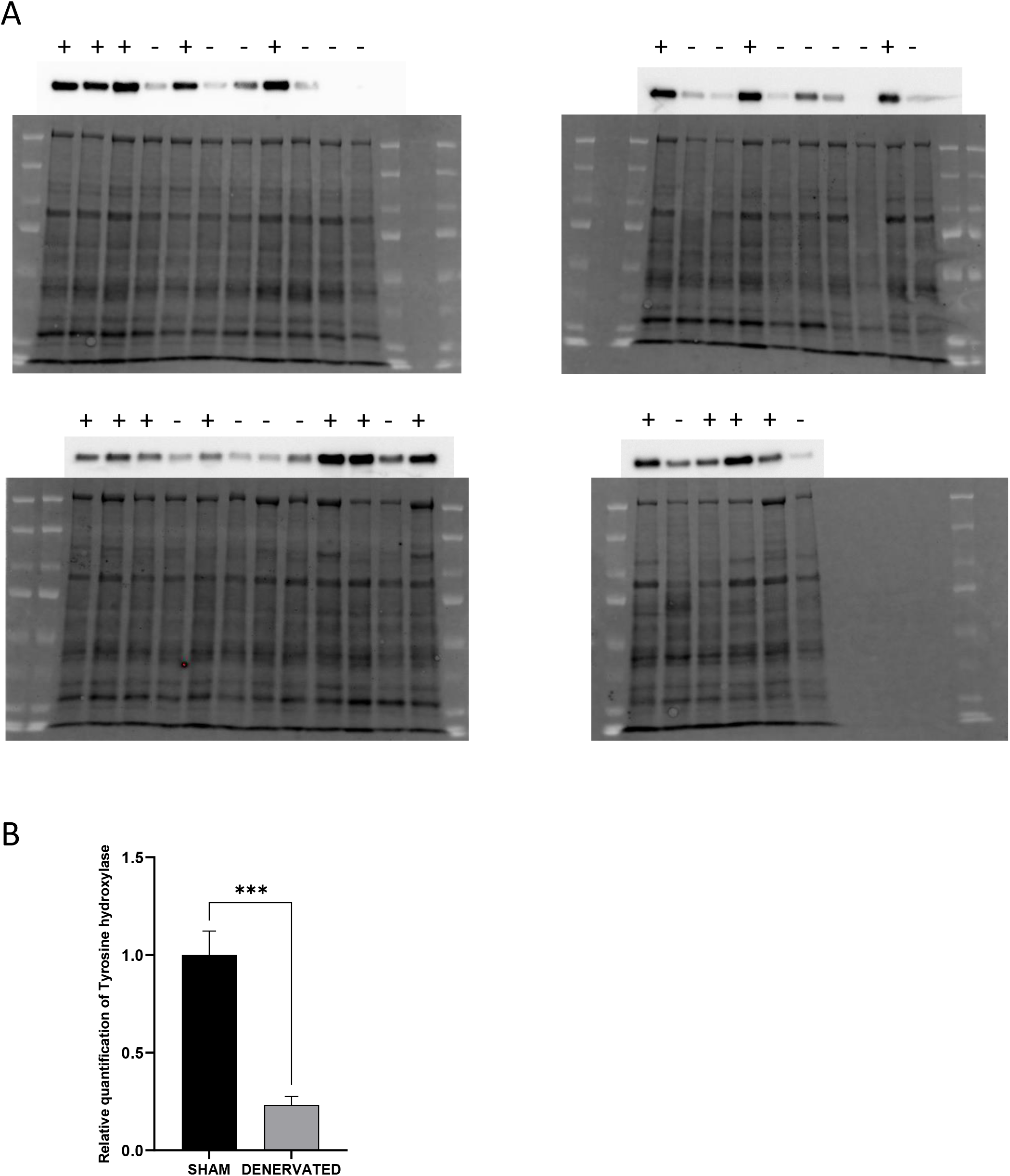
Confirmation of BAT denervation by quantification of tyrosine hydroxylase protein. **(A)** Western-blot with total protein (bottom of the pictures) and tyrosine hydroxylase (top of the pictures) extracted from BAT. Each lane represents a different sample, from sham mice (+) or denervated mice (-). **(B)** Relative quantification of tyrosine hydroxylase protein in the BAT of sham and denervated mice. Quantity of tyrosine hydroxylase is relative to the quantity of total proteins. Error bars represent the SEM. ***p < 0.001 versus SHAM mice

**Supplementary figure 4:**
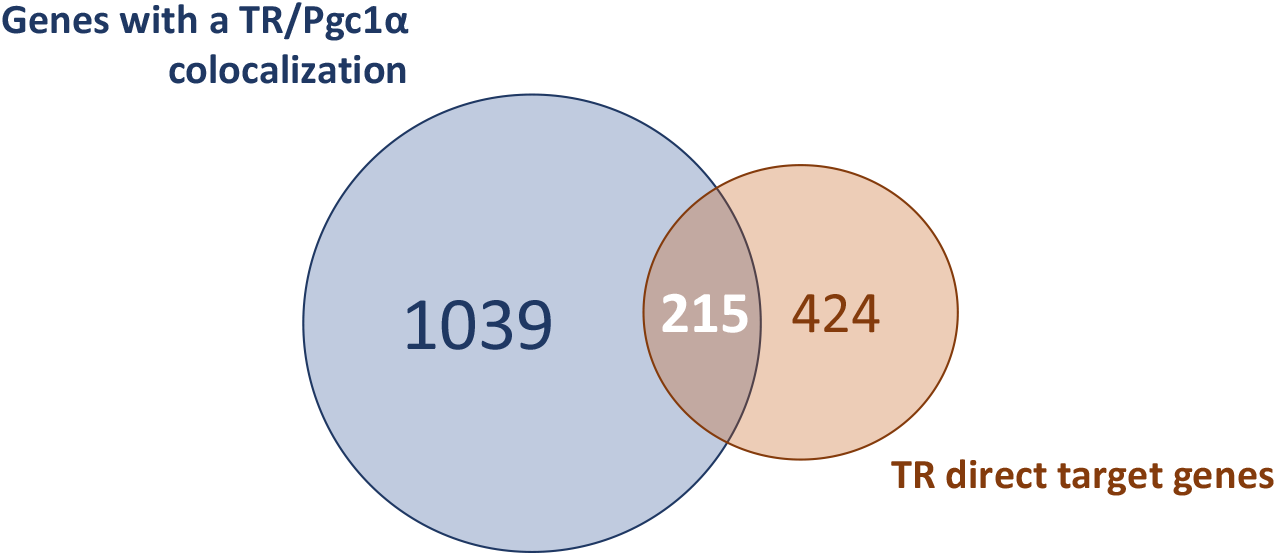
Comparison of TR direct target genes with the genes presenting a PGC1α binding site. Venn diagram designed to highlight TR direct target genes that include a PGC1α binding site.

**Supplementary figure 5:**
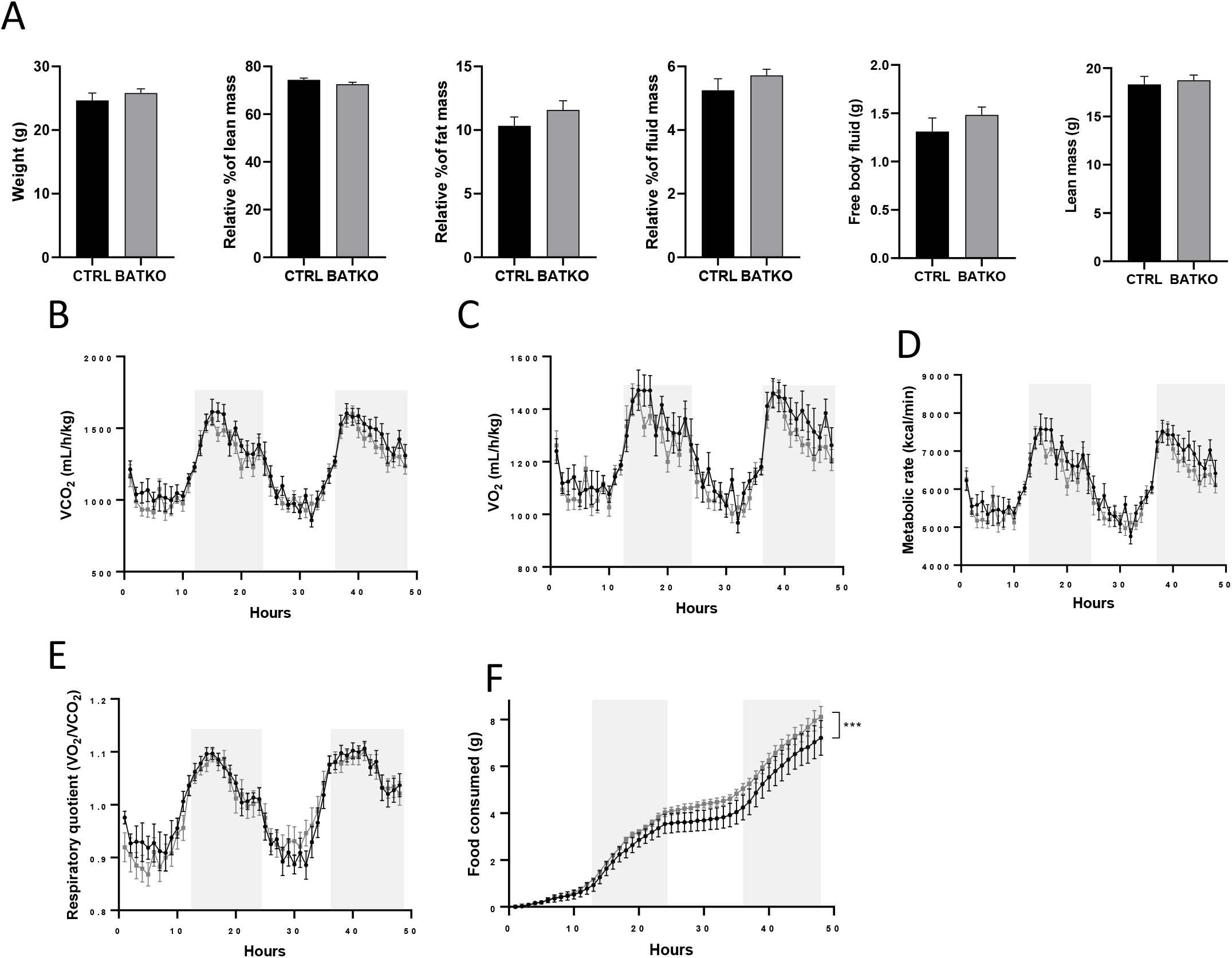
Body composition and indirect calorimetry in CTRL and BATKO mice. **(A)** Body composition of BATKO and CTRL mice measured by nuclear magnetic resonance (n=7-9/group). **(B)** VCO_2_(mL/h/kg), **(C)** VO_2_ (mL/h/kg), **(D)** Metabolic rate, **(E)** Respiratory quotient (ratio of VO_2_/VCO_2_), **(F)** cumulative food consumed by CTRL (black) and BATKO (grey) mice, as monitored during 48h with 12h light/dark cyles (dark phases are represented by grey areas). Error bars represent the SEM.

**Supplementary figure 6:**
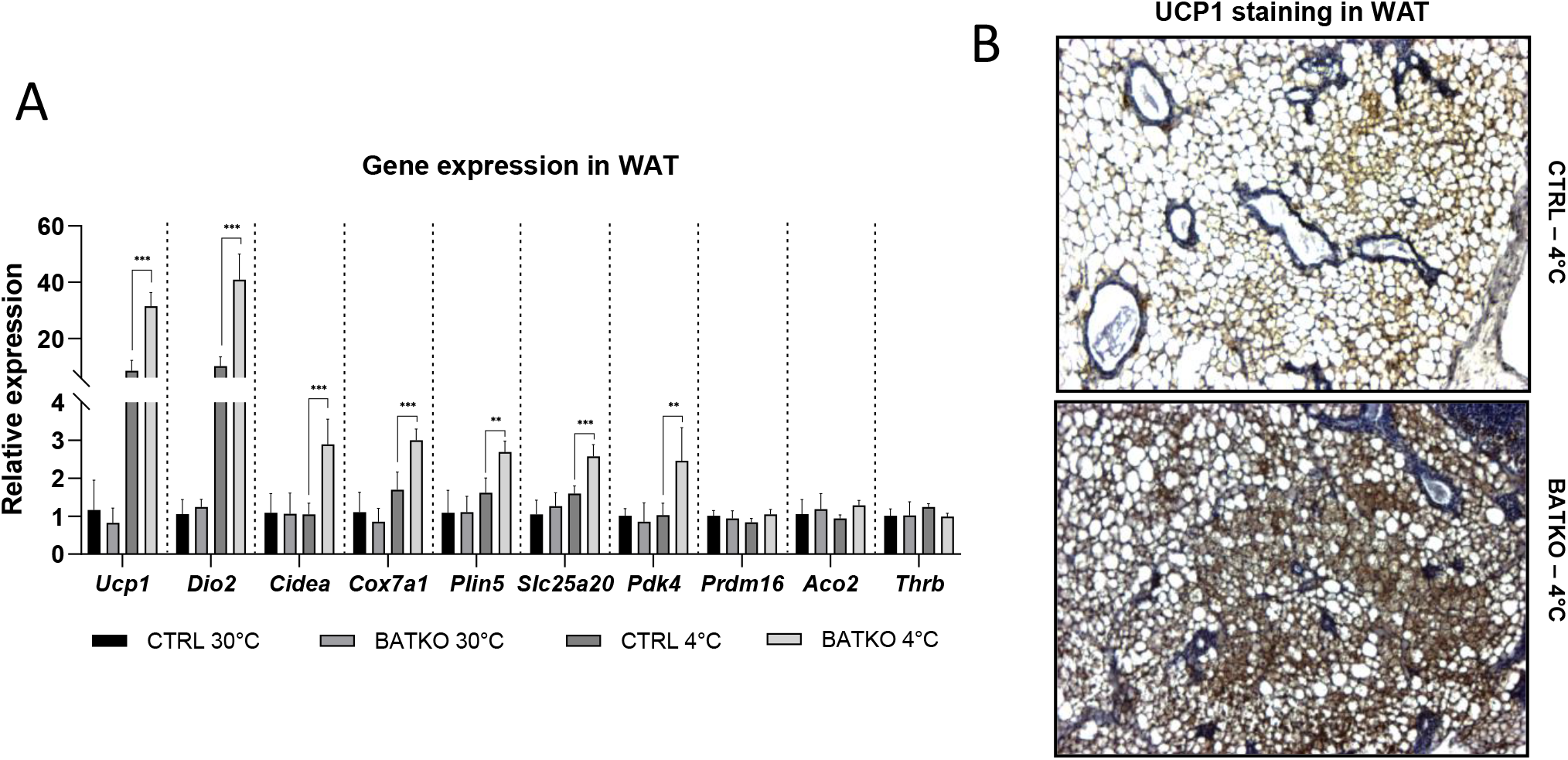
WAT browning is exacerbated in BATKO mice. **(A)** Relative expression of genes in the WAT (n=4-7/group) and **(B)** representative images of UCP1 staining in the WAT, in CTRL and BATKO mice after a 72h 4°C exposure (with *ad libitum* access to food). Browning was exacerbated in BATKO mice. *Thrb* was not affected in WAT. Statistical significance is shown for the CTRL 4°C *vs* BATKO 4°C group comparison.

**Supplementary figure 7:**
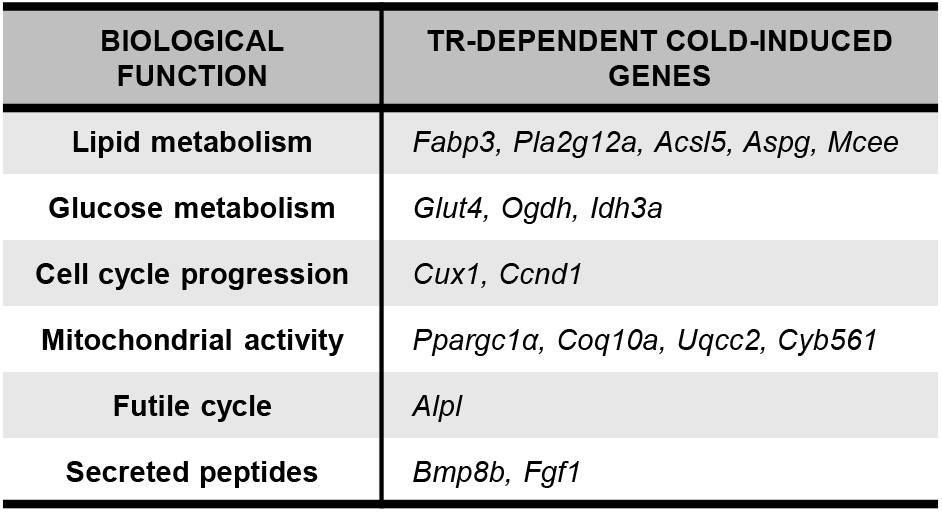
TR direct target genes are dysregulated during cold exposure in BATKO mice. Table of the genes inefficiently induced in BATKO mice at cold, restricted to the genes which are TR direct targets genes in BAT. These genes were manually attributed to their main biological function.

**Supplementary figure 8:**
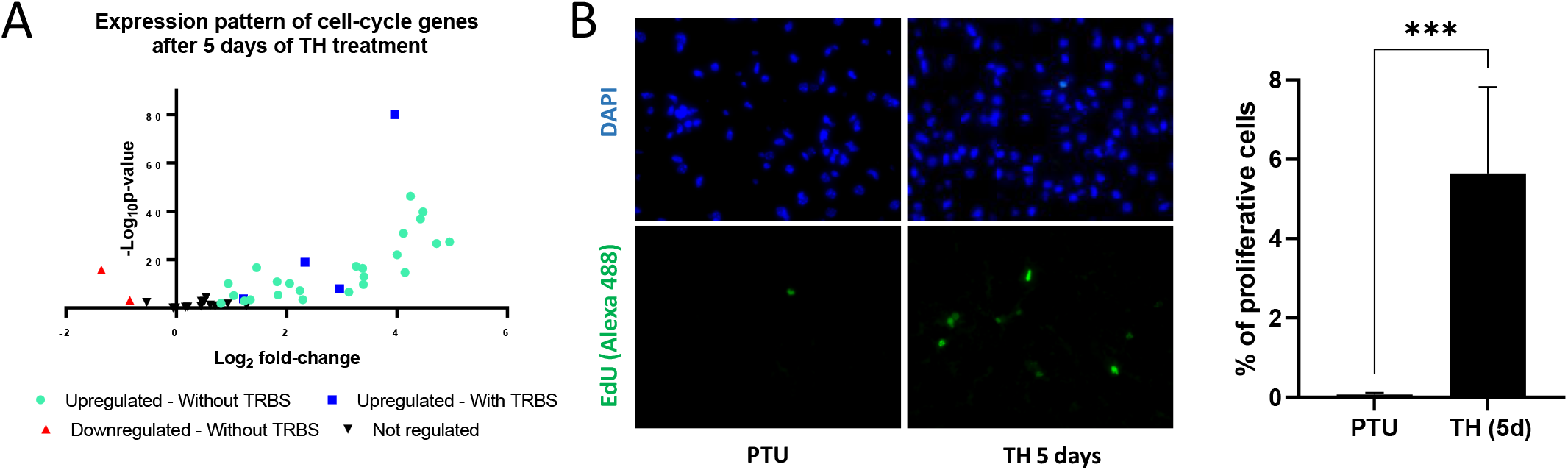
TH trigger BAT proliferation. **(A)** RNAseq expression pattern of cell-cycle genes(36) after 5 days of TH treatment in wild-type PTU-fed mice. Genes are divided into 4 groups, according to their regulation by TH treatment (n=4/group) and the presence of a TRBS within 30kb from their promoter. **(B)** Representative images (left) and quantification (right) of EdU-positive proliferative cells (green cells) in BAT from wild-type hypothyroid mice treated or not with TH during 5 days and co-injected with EdU at day 4. Nuclei are stained in blue with DAPI. Percentage of proliferative cells is the ratio of proliferative cells on the number of nuclei (n=5-6/group).

**Supplementary figure 9:**
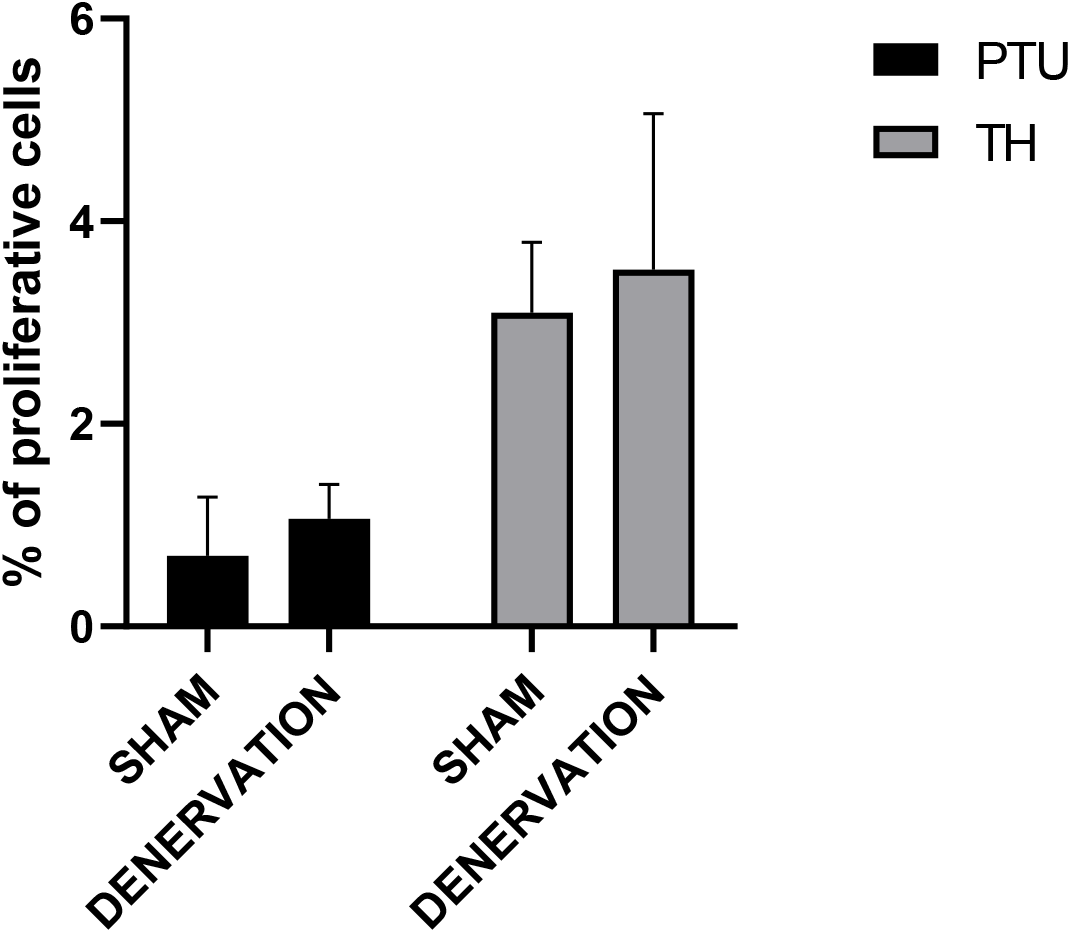
Effects of BAT denervation on T3-induced proliferation. Quantification of EdU positive cells in BAT from PTU-fed denervated or sham mice treated or not with TH during 5 days and co-injected with EdU at day 4. Percentage of proliferative cells is the ratio of proliferative cells on the number of nuclei (n=5/group). Error bars represent the SD.

## References

(1) Tabuchi, C.; Sul, H. S. Signaling Pathways Regulating Thermogenesis. Front Endocrinol (Lausanne) 2021, 12, 595020. https://doi.org/10.3389/fendo.2021.595020.

(2) Hankir, M. K.; Klingenspor, M. Brown Adipocyte Glucose Metabolism: A Heated Subject. EMBO Rep 2018, 19 (9), e46404. https://doi.org/10.15252/embr.201846404.

(3) Fenzl, A.; Kiefer, F. W. Brown Adipose Tissue and Thermogenesis. Horm Mol Biol Clin Investig 2014, 19 (1), 25–37. https://doi.org/10.1515/hmbci-2014-0022.

(4) Fonseca, T. L.; Werneck-De-Castro, J. P.; Castillo, M.; Bocco, B. M. L. C.; Fernandes, G. W.; McAninch, E. A.; Ignacio, D. L.; Moises, C. C. S.; Ferreira, A. R.; Gereben, B.; Bianco, A. C. Tissue-Specific Inactivation of Type 2 Deiodinase Reveals Multilevel Control of Fatty Acid Oxidation by Thyroid Hormone in the Mouse. Diabetes 2014, 63 (5), 1594–1604. https://doi.org/10.2337/db13-1768.

(5) Uldry, M.; Yang, W.; St-Pierre, J.; Lin, J.; Seale, P.; Spiegelman, B. M. Complementary Action of the PGC-1 Coactivators in Mitochondrial Biogenesis and Brown Fat Differentiation. Cell Metab 2006, 3 (5), 333–341. https://doi.org/10.1016/j.cmet.2006.04.002.

(6) Fukano, K.; Okamatsu-Ogura, Y.; Tsubota, A.; Nio-Kobayashi, J.; Kimura, K. Cold Exposure Induces Proliferation of Mature Brown Adipocyte in a SS3-Adrenergic Receptor-Mediated Pathway. PLoS One 2016, 11 (11), e0166579. https://doi.org/10.1371/journal.pone.0166579.

(7) Cao, W.; Daniel, K. W.; Robidoux, J.; Puigserver, P.; Medvedev, A. V.; Bai, X.; Floering, L. M.; Spiegelman, B. M.; Collins, S. P38 Mitogen-Activated Protein Kinase Is the Central Regulator of Cyclic AMP-Dependent Transcription of the Brown Fat Uncoupling Protein 1 Gene. Mol Cell Biol 2004, 24 (7), 3057–3067. https://doi.org/10.1128/MCB.24.7.3057-3067.2004.

(8) Silva, J. E.; Larsen, P. R. Potential of Brown Adipose Tissue Type II Thyroxine 5’-Deiodinase as a Local and Systemic Source of Triiodothyronine in Rats. J Clin Invest 1985, 76 (6), 2296–2305. https://doi.org/10.1172/JCI112239.

(9) Bianco, A. C.; Silva, J. E. Intracellular Conversion of Thyroxine to Triiodothyronine Is Required for the Optimal Thermogenic Function of Brown Adipose Tissue. J Clin Invest 1987, 79 (1), 295–300. https://doi.org/10.1172/JCI112798.

(10) Maushart, C. I.; Loeliger, R.; Gashi, G.; Christ-Crain, M.; Betz, M. J. Resolution of Hypothyroidism Restores Cold-Induced Thermogenesis in Humans. Thyroid 2019, 29 (4), 493–501. https://doi.org/10.1089/thy.2018.0436.

(11) Wolf, M.; Weigert, A.; Kreymann, G. Body Composition and Energy Expenditure in Thyroidectomized Patients during Short-Term Hypothyroidism and Thyrotropin-Suppressive Thyroxine Therapy. Eur J Endocrinol 1996, 134 (2), 168–173. https://doi.org/10.1530/eje.0.1340168.

(12) De Leo, S.; Lee, S. Y.; Braverman, L. E. Hyperthyroidism. Lancet 2016, 388 (10047), 906–918. https://doi.org/10.1016/S0140-6736(16)00278-6.

(13) Minakhina, S.; Bansal, S.; Zhang, A.; Brotherton, M.; Janodia, R.; De Oliveira, V.; Tadepalli, S.; Wondisford, F. E. A Direct Comparison of Thyroid Hormone Receptor Protein Levels in Mice Provides Unexpected Insights into Thyroid Hormone Action. Thyroid 2020, 30 (8), 1193–1204. https://doi.org/10.1089/thy.2019.0763.

(14) Flamant, F. Futures Challenges in Thyroid Hormone Signaling Research. Front Endocrinol (Lausanne) 2016, 7, 58. https://doi.org/10.3389/fendo.2016.00058.

(15) Bianco, A. C.; Silva, J. E. Cold Exposure Rapidly Induces Virtual Saturation of Brown Adipose Tissue Nuclear T3 Receptors. Am J Physiol 1988, 255 (4 Pt 1), E496–503. https://doi.org/10.1152/ajpendo.1988.255.4.E496.

(16) Schneider, M. J.; Fiering, S. N.; Pallud, S. E.; Parlow, A. F.; St. Germain, D. L.; Galton, V. A. Targeted Disruption of the Type 2 Selenodeiodinase Gene (DIO2) Results in a Phenotype of Pituitary Resistance to T4. Molecular Endocrinology 2001, 15 (12), 2137–2148. https://doi.org/10.1210/mend.15.12.0740.

(17) Ritter, M. J.; Amano, I.; Hollenberg, A. N. Thyroid Hormone Signaling and the Liver. Hepatology 2020, 72 (2), 742–752. https://doi.org/10.1002/hep.31296.

(18) Nicolaisen, T. S.; Klein, A. B.; Dmytriyeva, O.; Lund, J.; Ingerslev, L. R.; Fritzen, A. M.; Carl, C. S.; Lundsgaard, A.-M.; Frost, M.; Ma, T.; Schjerling, P.; Gerhart-Hines, Z.; Flamant, F.; Gauthier, K.; Larsen, S.; Richter, E. A.; Kiens, B.; Clemmensen, C. Thyroid Hormone Receptor α in Skeletal Muscle Is Essential for T3-Mediated Increase in Energy Expenditure. FASEB J 2020, 34 (11), 15480–15491. https://doi.org/10.1096/fj.202001258RR.

(19) Martínez-Sánchez, N.; Moreno-Navarrete, J. M.; Contreras, C.; Rial-Pensado, E.; Fernø, J.; Nogueiras, R.; Diéguez, C.; Fernández-Real, J.-M.; López, M. Thyroid Hormones Induce Browning of White Fat. J Endocrinol 2016, 232 (2), 351–362. https://doi.org/10.1530/JOE-16-0425.

(20) Medina-Gomez, G.; Calvo, R. M.; Obregon, M.-J. Thermogenic Effect of Triiodothyroacetic Acid at Low Doses in Rat Adipose Tissue without Adverse Side Effects in the Thyroid Axis. Am J Physiol Endocrinol Metab 2008, 294 (4), E688–697. https://doi.org/10.1152/ajpendo.00417.2007.

(21) López, M.; Varela, L.; Vázquez, M. J.; Rodríguez-Cuenca, S.; González, C. R.; Velagapudi, V. R.; Morgan, D. A.; Schoenmakers, E.; Agassandian, K.; Lage, R.; Martínez de Morentin, P. B.; Tovar, S.; Nogueiras, R.; Carling, D.; Lelliott, C.; Gallego, R.; Oresic, M.; Chatterjee, K.; Saha, A. K.; Rahmouni, K.; Diéguez, C.; Vidal-Puig, A. Hypothalamic AMPK and Fatty Acid Metabolism Mediate Thyroid Regulation of Energy Balance. Nat Med 2010, 16 (9), 1001–1008. https://doi.org/10.1038/nm.2207.

(22) Quignodon, L.; Vincent, S.; Winter, H.; Samarut, J.; Flamant, F. A Point Mutation in the Activation Function 2 Domain of Thyroid Hormone Receptor A1 Expressed after CRE-Mediated Recombination Partially Recapitulates Hypothyroidism. Molecular Endocrinology 2007, 21 (10), 2350–2360. https://doi.org/10.1210/me.2007-0176.

(23) Winter, H.; Rüttiger, L.; Müller, M.; Kuhn, S.; Brandt, N.; Zimmermann, U.; Hirt, B.; Bress, A.; Sausbier, M.; Conscience, A.; Flamant, F.; Tian, Y.; Zuo, J.; Pfister, M.; Ruth, P.; Löwenheim, H.; Samarut, J.; Engel, J.; Knipper, M. Deafness in TRβ Mutants Is Caused by Malformation of the Tectorial Membrane. J Neurosci 2009, 29 (8), 2581–2587. https://doi.org/10.1523/JNEUROSCI.3557-08.2009.

(24) Potter, G. B.; Zarach, J. M.; Sisk, J. M.; Thompson, C. C. The Thyroid Hormone-Regulated Corepressor Hairless Associates with Histone Deacetylases in Neonatal Rat Brain. Mol Endocrinol 2002, 16 (11), 2547–2560. https://doi.org/10.1210/me.2002-0115.

(25) Engelhard, A.; Christiano, A. M. The Hairless Promoter Is Differentially Regulated by Thyroid Hormone in Keratinocytes and Neuroblastoma Cells. Exp Dermatol 2004, 13 (4), 257–264. https://doi.org/10.1111/j.0906-6705.2004.00175.x.

(26) Biagi, C. A. O.; Cury, S. S.; Alves, C. P.; Rabhi, N.; Silva, W. A.; Farmer, S. R.; Carvalho, R. F.; Batista, M. L. Multidimensional Single-Nuclei RNA-Seq Reconstruction of Adipose Tissue Reveals Adipocyte Plasticity Underlying Thermogenic Response. Cells 2021, 10 (11), 3073. https://doi.org/10.3390/cells10113073.

(27) Chatonnet, F.; Guyot, R.; Benoît, G.; Flamant, F. Genome-Wide Analysis of Thyroid Hormone Receptors Shared and Specific Functions in Neural Cells. Proc Natl Acad Sci U S A 2013, 110 (8), E766–775. https://doi.org/10.1073/pnas.1210626110.

(28) Hirose, K.; Payumo, A. Y.; Cutie, S.; Hoang, A.; Zhang, H.; Guyot, R.; Lunn, D.; Bigley, R. B.; Yu, H.; Wang, J.; Smith, M.; Gillett, E.; Muroy, S. E.; Schmid, T.; Wilson, E.; Field, K. A.; Reeder, D. M.; Maden, M.; Yartsev, M. M.; Wolfgang, M. J.; Grützner, F.; Scanlan, T. S.; Szweda, L. I.; Buffenstein, R.; Hu, G.; Flamant, F.; Olgin, J. E.; Huang, G. N. Evidence for Hormonal Control of Heart Regenerative Capacity during Endothermy Acquisition. Science 2019, 364 (6436), 184–188. https://doi.org/10.1126/science.aar2038.

(29) Richard, S.; Guyot, R.; Rey-Millet, M.; Prieux, M.; Markossian, S.; Aubert, D.; Flamant, F. A Pivotal Genetic Program Controlled by Thyroid Hormone during the Maturation of GABAergic Neurons. iScience 2020, 23 (3), 100899. https://doi.org/10.1016/j.isci.2020.100899.

(30) Martinez de Mena, R.; Scanlan, T. S.; Obregon, M.-J. The T3 Receptor Beta1 Isoform Regulates UCP1 and D2 Deiodinase in Rat Brown Adipocytes. Endocrinology 2010, 151 (10), 5074–5083. https://doi.org/10.1210/en.2010-0533.

(31) Yuan, C.; Nguyen, P.; Baxter, J. D.; Webb, P. Distinct Ligand-Dependent and Independent Modes of Thyroid Hormone Receptor (TR)/PGC-1α Interaction. J Steroid Biochem Mol Biol 2013, 133, 58–65. https://doi.org/10.1016/j.jsbmb.2012.09.001.

(32) Chang, J. S.; Ghosh, S.; Newman, S.; Salbaum, J. M. A Map of the PGC-1α- and NT-PGC-1α-Regulated Transcriptional Network in Brown Adipose Tissue. Sci Rep 2018, 8 (1), 7876. https://doi.org/10.1038/s41598-018-26244-4.

(33) Reitman, M. L. Of Mice and Men – Environmental Temperature, Body Temperature, and Treatment of Obesity. FEBS Letters 2018, 592 (12), 2098–2107. https://doi.org/10.1002/1873-3468.13070.

(34) Love, M. I.; Huber, W.; Anders, S. Moderated Estimation of Fold Change and Dispersion for RNA-Seq Data with DESeq2. Genome Biol 2014, 15 (12), 550. https://doi.org/10.1186/s13059-014-0550-8.

(35) Jeong, J. H.; Chang, J. S.; Jo, Y.-H. Intracellular Glycolysis in Brown Adipose Tissue Is Essential for Optogenetically Induced Nonshivering Thermogenesis in Mice. Sci Rep 2018, 8 (1), 6672. https://doi.org/10.1038/s41598-018-25265-3.

(36) Whitfield, M. L.; Sherlock, G.; Saldanha, A. J.; Murray, J. I.; Ball, C. A.; Alexander, K. E.; Matese, J. C.; Perou, C. M.; Hurt, M. M.; Brown, P. O.; Botstein, D. Identification of Genes Periodically Expressed in the Human Cell Cycle and Their Expression in Tumors. Mol Biol Cell 2002, 13 (6), 1977–2000. https://doi.org/10.1091/mbc.02-02-0030.

(37) Hall, J. A.; Ribich, S.; Christoffolete, M. A.; Simovic, G.; Correa-Medina, M.; Patti, M. E.; Bianco, A. C. Absence of Thyroid Hormone Activation during Development Underlies a Permanent Defect in Adaptive Thermogenesis. Endocrinology 2010, 151 (9), 4573–4582. https://doi.org/10.1210/en.2010-0511.

(38) Wulf, A.; Harneit, A.; Kröger, M.; Kebenko, M.; Wetzel, M. G.; Weitzel, J. M. T3-Mediated Expression of PGC-1alpha via a Far Upstream Located Thyroid Hormone Response Element. Mol Cell Endocrinol 2008, 287 (1–2), 90–95. https://doi.org/10.1016/j.mce.2008.01.017.

(39) Cannon, B.; Nedergaard, J. Brown Adipose Tissue: Function and Physiological Significance. Physiol Rev 2004, 84 (1), 277–359. https://doi.org/10.1152/physrev.00015.2003.

(40) Silvestri, E.; Senese, R.; De Matteis, R.; Cioffi, F.; Moreno, M.; Lanni, A.; Gentile, A.; Busiello, R. A.; Salzano, A. M.; Scaloni, A.; de Lange, P.; Goglia, F.; Lombardi, A. Absence of Uncoupling Protein 3 at Thermoneutrality Influences Brown Adipose Tissue Mitochondrial Functionality in Mice. FASEB J 2020, 34 (11), 15146–15163. https://doi.org/10.1096/fj.202000995R.

(41) Adams, A. C.; Astapova, I.; Fisher, F. M.; Badman, M. K.; Kurgansky, K. E.; Flier, J. S.; Hollenberg, A. N.; Maratos-Flier, E. Thyroid Hormone Regulates Hepatic Expression of Fibroblast Growth Factor 21 in a PPARalpha-Dependent Manner. J Biol Chem 2010, 285 (19), 14078–14082. https://doi.org/10.1074/jbc.C110.107375.

(42) Sáenz de Urturi, D.; Buqué, X.; Porteiro, B.; Folgueira, C.; Mora, A.; Delgado, T. C.; Prieto-Fernández, E.; Olaizola, P.; Gómez-Santos, B.; Apodaka-Biguri, M.; González-Romero, F.; Nieva-Zuluaga, A.; Ruiz de Gauna, M.; Goikoetxea-Usandizaga, N.; García-Rodríguez, J. L.; Gutierrez de Juan, V.; Aurrekoetxea, I.; Montalvo-Romeral, V.; Novoa, E. M.; Martín-Guerrero, I.; Varela-Rey, M.; Bhanot, S.; Lee, R.; Banales, J. M.; Syn, W.-K.; Sabio, G.; Martínez-Chantar, M. L.; Nogueiras, R.; Aspichueta, P. Methionine Adenosyltransferase 1a Antisense Oligonucleotides Activate the Liver-Brown Adipose Tissue Axis Preventing Obesity and Associated Hepatosteatosis. Nat Commun 2022, 13 (1), 1096. https://doi.org/10.1038/s41467-022-28749-z.

(43) de Jesus, L. A.; Carvalho, S. D.; Ribeiro, M. O.; Schneider, M.; Kim, S.-W.; Harney, J. W.; Larsen, P. R.; Bianco, A. C. The Type 2 Iodothyronine Deiodinase Is Essential for Adaptive Thermogenesis in Brown Adipose Tissue. J. Clin. Invest. 2001, 108 (9), 1379–1385. https://doi.org/10.1172/JCI200113803.

(44) Christoffolete, M. A.; Linardi, C. C. G.; de Jesus, L.; Ebina, K. N.; Carvalho, S. D.; Ribeiro, M. O.; Rabelo, R.; Curcio, C.; Martins, L.; Kimura, E. T.; Bianco, A. C. Mice with Targeted Disruption of the Dio2 Gene Have Cold-Induced Overexpression of the Uncoupling Protein 1 Gene but Fail to Increase Brown Adipose Tissue Lipogenesis and Adaptive Thermogenesis. Diabetes 2004, 53 (3), 577–584. https://doi.org/10.2337/diabetes.53.3.577.

(45) Oppenheimer, J. H.; Schwartz, H. L.; Lane, J. T.; Thompson, M. P. Functional Relationship of Thyroid Hormone-Induced Lipogenesis, Lipolysis, and Thermogenesis in the Rat. J. Clin. Invest. 1991, 87 (1), 125–132. https://doi.org/10.1172/JCI114961.

(46) Sun, Y.; Rahbani, J. F.; Jedrychowski, M. P.; Riley, C. L.; Vidoni, S.; Bogoslavski, D.; Hu, B.; Dumesic, P. A.; Zeng, X.; Wang, A. B.; Knudsen, N. H.; Kim, C. R.; Marasciullo, A.; Millán, J. L.; Chouchani, E. T.; Kazak, L.; Spiegelman, B. M. Mitochondrial TNAP Controls Thermogenesis by Hydrolysis of Phosphocreatine. Nature 2021, 593 (7860), 580–585. https://doi.org/10.1038/s41586-021-03533-z.

(47) Liu, S.; Shen, S.; Yan, Y.; Sun, C.; Lu, Z.; Feng, H.; Ma, Y.; Tang, Z.; Yu, J.; Wu, Y.; Gereben, B.; Mohácsik, P.; Fekete, C.; Feng, X.; Yuan, F.; Guo, F.; Hu, C.; Shao, M.; Gao, X.; Zhao, L.; Li, Y.; Jiang, J.; Ying, H. Triiodothyronine (T3) Promotes Brown Fat Hyperplasia via Thyroid Hormone Receptor α Mediated Adipocyte Progenitor Cell Proliferation. Nat Commun 2022, 13, 3394. https://doi.org/10.1038/s41467-022-31154-1.

(48) Pascual, A.; Aranda, A. Thyroid Hormone Receptors, Cell Growth and Differentiation. Biochimica et Biophysica Acta (BBA) - General Subjects 2013, 1830 (7), 3908–3916. https://doi.org/10.1016/j.bbagen.2012.03.012.

(49) Škop, V.; Guo, J.; Liu, N.; Xiao, C.; Hall, K. D.; Gavrilova, O.; Reitman, M. L. Mouse Thermoregulation: Introducing the Concept of the Thermoneutral Point. Cell Rep 2020, 31 (2), 107501. https://doi.org/10.1016/j.celrep.2020.03.065.

(50) Challa, T. D.; Dapito, D. H.; Kulenkampff, E.; Kiehlmann, E.; Moser, C.; Straub, L.; Sun, W.; Wolfrum, C. A Genetic Model to Study the Contribution of Brown and Brite Adipocytes to Metabolism. Cell Rep 2020, 30 (10), 3424–3433.e4. https://doi.org/10.1016/j.celrep.2020.02.055.

(51) Castillo, M.; Hall, J. A.; Correa-Medina, M.; Ueta, C.; Kang, H. W.; Cohen, D. E.; Bianco, A. C. Disruption of Thyroid Hormone Activation in Type 2 Deiodinase Knockout Mice Causes Obesity with Glucose Intolerance and Liver Steatosis Only at Thermoneutrality. Diabetes 2011, 60 (4), 1082–1089. https://doi.org/10.2337/db10-0758.

(52) Rosenwald, M.; Perdikari, A.; Rülicke, T.; Wolfrum, C. Bi-Directional Interconversion of Brite and White Adipocytes. Nat Cell Biol 2013, 15 (6), 659–667. https://doi.org/10.1038/ncb2740.

(53) Madisen, L.; Zwingman, T. A.; Sunkin, S. M.; Oh, S. W.; Zariwala, H. A.; Gu, H.; Ng, L. L.; Palmiter, R. D.; Hawrylycz, M. J.; Jones, A. R.; Lein, E. S.; Zeng, H. A Robust and High-Throughput Cre Reporting and Characterization System for the Whole Mouse Brain. Nat Neurosci 2010, 13 (1), 133–140. https://doi.org/10.1038/nn.2467.

(54) Weiss, R. E.; Murata, Y.; Cua, K.; Hayashi, Y.; Seo, H.; Refetoff, S. Thyroid Hormone Action on Liver, Heart, and Energy Expenditure in Thyroid Hormone Receptor Beta-Deficient Mice. Endocrinology 1998, 139 (12), 4945–4952. https://doi.org/10.1210/endo.139.12.6412.

(55) Weir, J. B. New Methods for Calculating Metabolic Rate with Special Reference to Protein Metabolism. 1949. Nutrition 1990, 6 (3), 213–221.

(56) Nitta, A.; Furukawa, Y.; Hayashi, K.; Hiramatsu, M.; Kameyama, T.; Hasegawa, T.; Nabeshima, T. Denervation of Dopaminergic Neurons with 6-Hydroxydopamine Increases Nerve Growth Factor Content in Rat Brain. Neurosci Lett 1992, 144 (1–2), 152–156. https://doi.org/10.1016/0304-3940(92)90738-s.

(57) Bookout, A. L.; Mangelsdorf, D. J. Quantitative Real-Time PCR Protocol for Analysis of Nuclear Receptor Signaling Pathways. Nucl Recept Signal 2003, 1, e012. https://doi.org/10.1621/nrs.01012.

(58) Guyot, R.; Chatonnet, F.; Gillet, B.; Hughes, S.; Flamant, F. Toxicogenomic Analysis of the Ability of Brominated Flame Retardants TBBPA and BDE-209 to Disrupt Thyroid Hormone Signaling in Neural Cells. Toxicology 2014, 325, 125–132. https://doi.org/10.1016/j.tox.2014.08.007.

(59) Ge, X. IDEP Web Application for RNA-Seq Data Analysis. Methods Mol Biol 2021, 2284, 417–443. https://doi.org/10.1007/978-1-0716-1307-8_22.

(60) Gil-Ibañez, P.; Morte, B.; Bernal, J. Role of Thyroid Hormone Receptor Subtypes α and β on Gene Expression in the Cerebral Cortex and Striatum of Postnatal Mice. Endocrinology 2013, 154 (5), 1940–1947. https://doi.org/10.1210/en.2012-2189.

